# Manipulable object and human contact: preferences and modulation of emotional states in weaned piglets

**DOI:** 10.1101/2020.06.29.177162

**Authors:** Avelyne S. Villain, Mathilde Lanthony, Carole Guérin, Camille Noûs, Céline Tallet

## Abstract

Enriching the life of farm animals is an obligation in intensive farming conditions. In pigs, manipulable materials are mandatory when no bedding is available. Like manipulable objects, positive human interactions might be considered as enrichment, as they provide the animals occasions to interact, increase their activity and lead to positive emotional states. In this study, we investigated how weaned piglets perceived a manipulable object, and a familiar human. After a similar familiarization to both stimuli, twenty-four weaned piglets were tested for a potential preference for one of the stimuli and submitted to isolation/reunion tests to evaluate the emotional value of the stimuli. We hypothesized that being reunited with a stimulus would attenuate the stress of social isolation and promote positive behaviors, and even more that the stimulus has a positive emotional value for piglets. Although our behavioural data did not allow to show a preference for one of the stimuli, piglets approached more often the human and were observed laying down only near the human. Using behavioural and bioacoustic data, we showed that reunion with the human decreased more the time spent in an attentive state and mobility of piglets than reunion with the object, and isolation. Vocalizations differed between reunions with the object and the human, and were different from vocalizations during isolation. The human presence led to higher frequency range, more noisy and shorter grunts. Finally, both stimuli decreased the isolation stress of piglets, and piglets seemed to be in a more positive emotional state with the human compared to the object. It confirms the potential need for positive human interactions to be used as pseudo-social enrichment in pigs.

## 2 Introduction

The intensive production system of animal products sometimes implies large densities of farm animals and can lead to deleterious behaviors and decrease the physical or mental health of animals, i.e. their welfare. Animal welfare covers, among others, the importance of the animal’s ability to keep control of mental and body stability in different environmental conditions (Broom 2011). Improving animal welfare is both reducing negatively perceived contexts as well as increasing positively perceived contexts and species-specific behaviors (Peterson, Simonsen, and Lawson 1995; Weerd and Day 2009). The pressure from citizens, consumers and animal welfare organizations has been growing regarding animal rights, leading to changes in the legislation. For example, the provision of manipulative materials to pigs of all ages is mandatory in the European Union since January 2013 (Council Directive 2008/120/EC 2008), materials named as ‘environmental enrichments’. Environmental enrichments are defined as materials susceptible to improve the biological functioning of captive animals (Newberry 1995) and should stimulate species-typical animals’ sensory systems, cognitive capacities and behaviours (Wells 2009). For instance, for pigs, enrichments materials should be edible, chewable, investigable, and manipulable (reviewed in Godyń, Nowicki, and Herbut 2019). Moreover, enrichment materials should be provided in such a way that they are of sustainable attraction for pigs, and should be accessible for oral manipulation, and provided in sufficient amount (Newberry 1995; Godyń, Nowicki, and Herbut 2019). Enrichments effects are generally tested using behavioural and physiological paradigms (Nannoni et al. 2016) and are classified as optimal (if they meet all of the above-mentioned criteria), suboptimal (if meet most of the criteria and should be combined with others) or marginal (they do not fulfill the animal need and should be used with other) (Godyń, Nowicki, and Herbut, 2019).

In the particular case of pigs, abnormal patterns of behavior (stereotypies, belly nosing, ear and tail biting) may arise at several stages of their development if they are prevented from any enrichment (Prunier et al. 2020). Enrichments have the potential to reduce these abnormal behaviours and increase positive behaviors like play (Lykhach et al. 2020; Luo et al. 2020). Although straw bedding is one optimal enrichment according to several literature references (reviewed in Godyń, Nowicki, and Herbut 2019), it is also non applicable for many farms using slatted floor systems. Thus manipulative materials have been developed and are used in farms in the form of ropes, hanging balls, wood, pipes or different commercial toys.

Besides those enrichment materials, one may wonder if enrichment may be provided by other stimuli in the environment of farm animals. Pigs being social animals, social enrichment is sometimes used for lactating piglets, by allowing different litters to interact. This enrichment enhances play and decreases aggression at weaning (Salazar et al. 2018). Another relational partner of pigs is their caregiver. Human interactions seems to correspond well to the definition of enrichment, i.e. they provide occasions of social contact with another animal (stimulates biological functioning), and stimulate all sensory systems of the animals. Humans, notably through their clothes and boots, are chewable, investigable, and manipulable. Many positive outcomes of positive human interactions have been shown. Farm animals may be tamed by humans providing regular positive additional contacts, leading to the expression of positive emotions (Tallet et al. 2018). Humans_may consequently be associated with positive outcomes as measured by a decrease of heart rate (Schmied et al. 2008; Tallet et al. 2014), modification of heart rate variability and ear postures (Coulon et al. 2015), EEG (Rault et al. 2019). Humans can also acquire reassuring properties in situations of social isolation (Rushen et al. 2001; Tallet, Veissier, and Boivin 2005). They may even induce behavioural reaction similar to those toward social partners (Brajon, Laforest, Bergeron, et al. 2015). Regular positive human contacts may even lead to improved welfare through positive cognitive bias (Brajon, Laforest, Schmitt, et al. 2015). In addition, pigs raised in a poor environment may develop more interest toward a familiar human than pigs raised in an enriched environment (Tallet et al. 2013), leading the author of the study to hypothesize a familiar human may be perceived as an enrichment. To our knowledge, no comparison exists on the effect of object enrichment and pseudo-social enrichment via human interactions. This would provide new insight into enrichment practices in pig breeding, for improving their welfare.

In this study, we developed a paradigm to test the perception piglets may have toward two stimuli: an inanimate object that could be used as enrichment, and a familiar human. The aim of this study was thus to elucidate the specific value that a familiar human may have compared to an inanimate object of enrichment. Twenty-four weaned piglets were previously similarly familiarized to both an object and a human. We first evaluated the potential preference for one of the stimuli. Then we evaluated the emotional value of the stimuli through isolation/reunion tests. We hypothesized that being reunited with a stimulus would attenuate the stress of social isolation and promote positive behaviors, and even more that the stimulus has a positive emotional value for piglets. We used behavioural but also bioacoustics data known to be relevant in comparing emotional states of pigs (Friel et al. 2019; Leliveld et al. 2017; Villain et al. 2020). Additionally, we tested if the level of attraction toward the stimuli could predict vocal expression in presence of each stimuli.

## 3 Material and methods

### 3.1 Ethical note

The experiment was held in France at the experimental farm UE3P, in Saint-Gilles, FRANCE. Experiments were performed under the authorization no. APAFIS#17071-2018101016045373_V3 from the French Ministry of Higher Education, Research and Innovation, received after evaluation of the regional ethic committee (Comité Rennais d’Ethique en Experimentation Animale); and were in agreement with the French and European legislation regarding experiments on animals.

### 3.2 Animals and experimental conditions

Twenty-four healthy weaned female piglets (Landrace/Large white dams inseminated with Piétrain semen) were involved in total. Piglets were weaned at 28 days of age. Eight groups of three sister piglets from eight different sows were selected at weaning according to their weight (the weight was balanced between and within the groups, 9.05± 0.66 kg on average). Groups were reared in the same rearing room in 115 x 132 cm pens, with slatted floor, isolated from each other by 1.5m high plastic panels. Piglets were fed *ad libitum* and had continuous access to a water trough. Each pen was provided with a steel chain as enrichment. The piglets were involved in the experiment from 28 to 57 days of age.

For different phases of the experiment, we also used an experimental room. The experimental room was located in the same building as the rearing room, 15 m away, and was a 270 cm x 270 cm soundproof room. It was warmed up by an electric heater. The entrance door was equipped with a hatch for the piglets. The transportation from the rearing room to the experimental room was done by the usual caretakers in closed trolleys. We used visually isolated trolleys to transport either the group of three piglets together (L120 x W80 x H80 cm) or one piglet at once (L80 x W50 x H80 cm).

### 3.3 Human and object familiarization

All the piglets were familiarized alternatively with two stimuli: an experimenter, referred as ‘Human’ in the rest of the article (always the same, a 1.65 m high woman dressed with a blue overall) and a 5L-plastic can (L20 x W10 x H30 cm) filled with water from which hung three pipe pieces tied with a thin rope, so that the three piglets could chew it all together, referred as ‘Object’ in the rest of the article. Familiarizations started at 28 days of age and ended at 53 days of age. Familiarizations were divided into two phases for each stimulus: eight sessions in the home pen (from days 29 to35), and eight sessions in the experimental room (from days 36 to 43 and from days 49 to 53). In the home pen, piglets received 10 minutes sessions twice a day for each stimulus along four days. During the same week, all groups were alternatively transported to the empty experimental room and remained there for 10 minutes, once a day for 5 days to get familiarized. After this familiarization, piglets received 10 minute sessions in the experimental room once a day for each stimulus, along nine days. The same procedure was used for each group of three pen mates, as follow:

- Object: the experimenter came before the grid of the pen holding the object and stood still and quiet for 30 seconds. Then, she entered the pen to tie the object to the grid opposite to the entrance with a small rope and went out. From the moment she went out, the object was left for 10 min in the pen. Then the experimenter removed the object.
- Human: the experimenter came in front of the grid of the pen and stood still, holding a stool, and quiet for 30 seconds. She then entered to sit on the stool in the pen, close to the grid at the opposite of the entrance. During the first session of human familiarization, the experimenter sat still without moving for 10 minutes before going out, in order not to afraid the piglets. For the other familiarization sessions, she engaged in interactions with each piglet (similar protocol as in Tallet et al. 2014): she hold out the hand towards the piglet; if the piglet did not move away, she tried to touch it; if the piglet accepted being touched, she softly stroked it along the body with the palm of her hand; and once it accepted being stroked, she scratched it along the body with her fingers. Scratching consisted in rubbing the skin of the piglets with the finger tips and applying more pressure than stroking. In addition, the handler spoke to the piglet with a soft voice. The experimenter focused on each piglet for two minutes first and alternatively focused her attention during the last four minutes.

The procedure of familiarization was similar in rearing pens and in the experimental room but the location of the stimulus changed: in rearing pens, the stimulus was attached to the opposite grid from the entrance of the pen. In the experimental room, the stimulus was placed in the center of the room.

### 3.4 Choice test

#### 3.4.1 Experimental procedure

At 47 and 49 days, the piglets were confronted twice with an individual Choice test between the familiar experimenter and the familiar object in order to evaluate the eventual preference for one of the stimuli. The test took place in the experimental room fitted in a V shaped apparatus (fig. 1A). Piglets had previously been familiarized to the room with the V shaped apparatus once for 5 minutes. The room was as much as possible made symmetrical, with false heater and camera, and a homogeneous ground surfacing. The day before, piglets where individually left five minutes in this room in order to get habituated to the room and being transported alone in a trolley. The day of the test, the piglets were brought individually at the entrance of the experimental room. Once in front of the experimental room, the hatch of the room and the first hatch of the trolley were opened for 30 s. Since the trolley had another grid hatch, the tested piglet could see into the experimental room without going out of the trolley. The human and the object were already in place at the back of the room (fig. 1A). The caretaker then opened the grid hatch and gently pushed the piglet into the room if it wouldn’t enter by itself after two seconds. The Choice test lasted five minutes. The experimenter called actively the piglet to come to her during the test. If the piglet got close, the experimenter petted it, similarly to the familiarization sessions. This test was done twice, on two days, swapping the sides of the stimuli between days in order to take into account eventual bias due to the laterality of the apparatus or the piglets.

**Figure 1:**
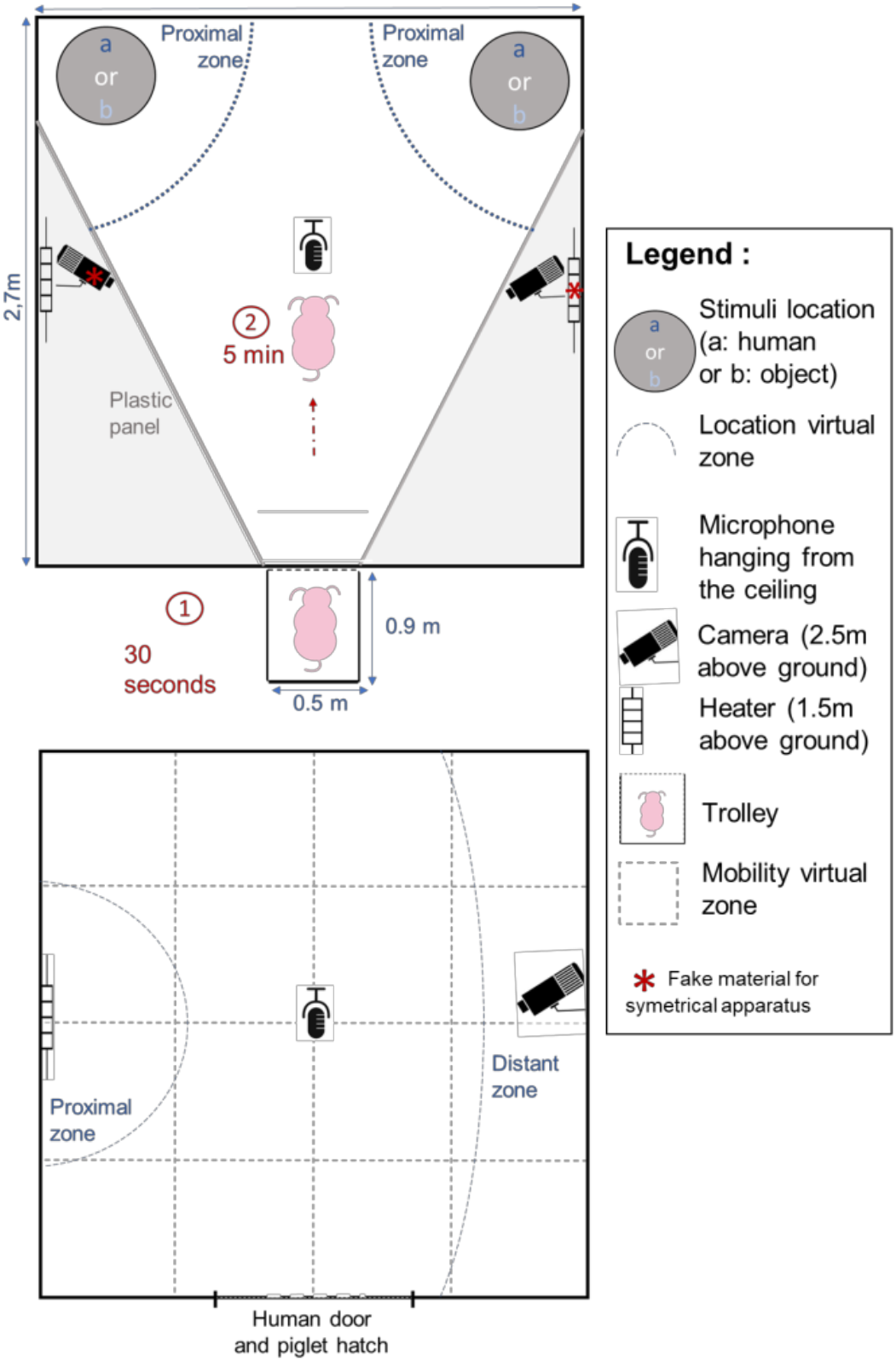
Design of the experimental apparatus for the Choice test (A) and the Isolation/Reunion test (B). For the Choice test between Object and Human (A), the piglets remained in the trolley for 30 seconds before entering the room where it was left for five minutes. For the Isolation/Reunion test (B), the piglet was left alone for five minutes (Isolation phase), and then remained for four minutes and 30 seconds (Reunion phase) either alone, with the familiar Object or with the familiar Human, depending days. Proximal zones: blue solid lines were drawn to identify the zones in which the piglet was considered close to the stimulus. The distant zone (B) was drawn to identify a zone where the piglet was considered distant from the stimulus. Virtual zones were drawn to monitor the location and mobility of the piglet in the room (grey dotted lines).

#### 3.4.2 Behavioural observations and analyses

The tests were video recorded by a camera (Bosh, Box 960H-CDD) and a recorder (Noldus Mpeg recorder 2.1., The Netherlands), linked to a LCD monitor (DELL 48 model 1907FPc) which allowed us to visualize the experimental room from an adjacent room. The location of the piglets was monitored directly during the tests and the other behaviors later from videos, both using the software The Observer XT 14.0 (Noldus^®^, The Netherlands). All behaviors recorded are indicated in table 1 and correspond to numbers: 2-7, 11, 14 (restricted to stimulus zone) and 16.

**Table 1:**
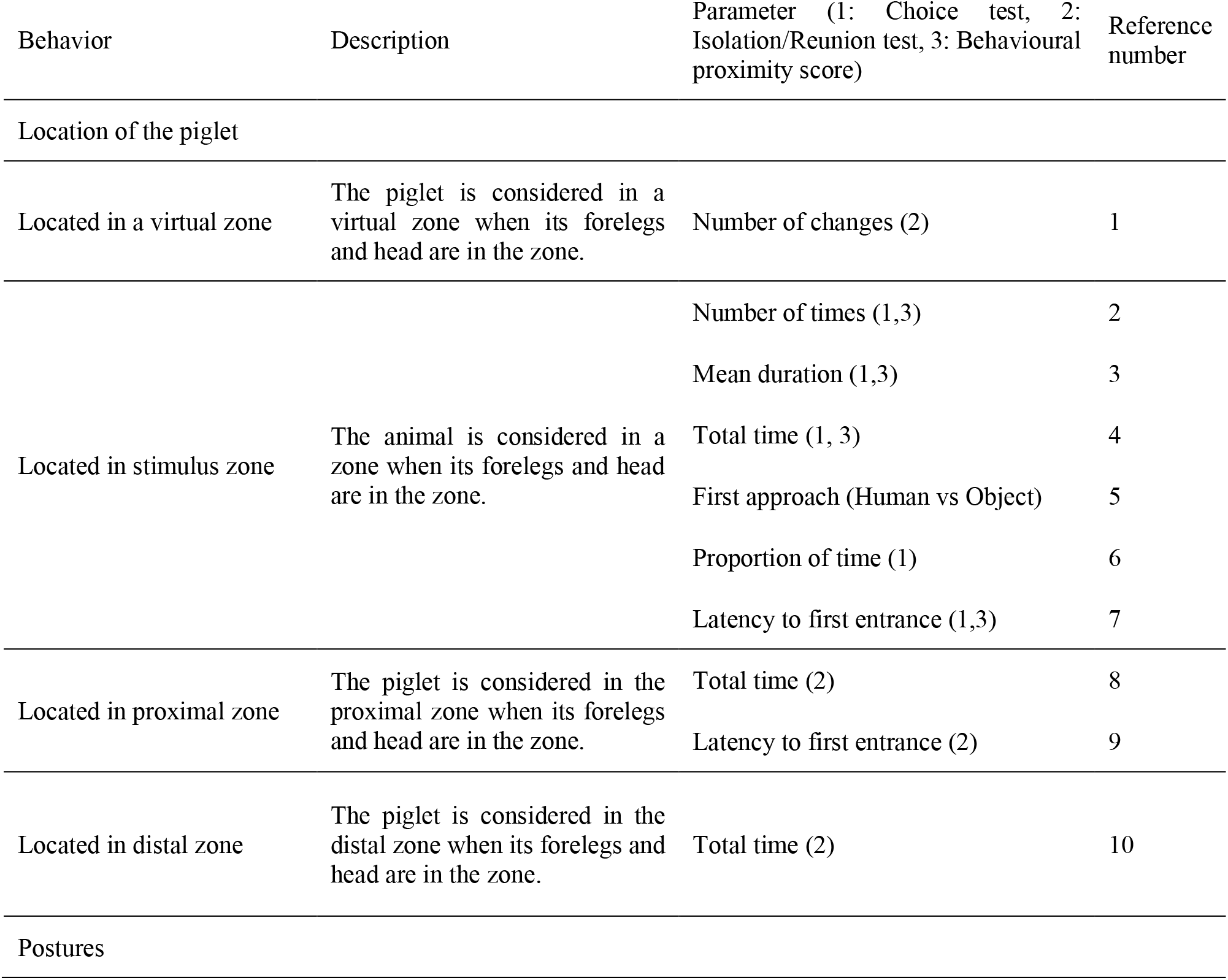
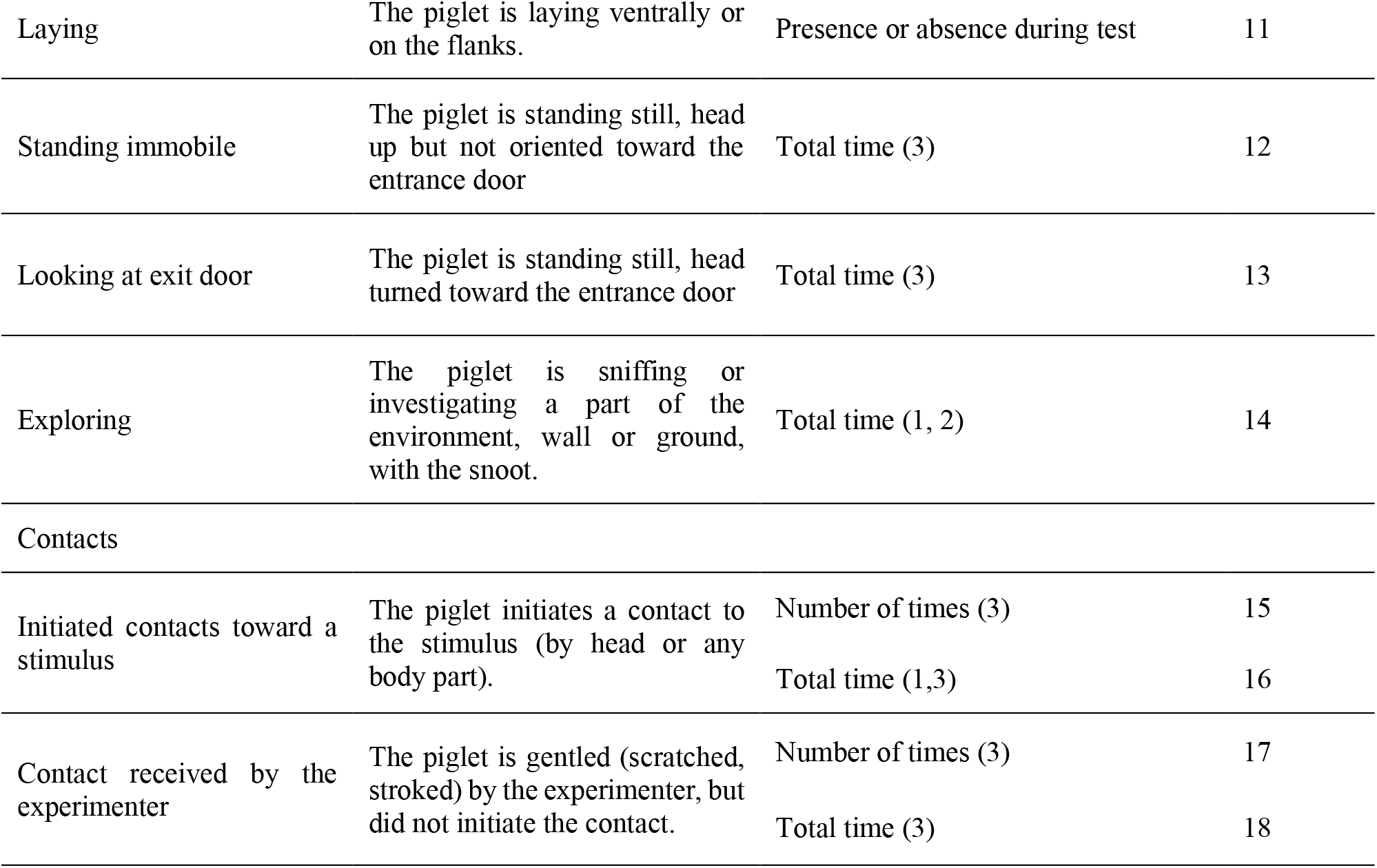
Ethogram used for the Choice test (1), Isolation/Reunion test (2) and behavioural proximity scores (3). Columns describe the name of the behavior, its description, the parameters calculated with it and for which test it was used. A number was assigned to each behavior for reference. The unit for timing is the second, except when it is labelled with a ‘*’ for which it is standardized per minute.

### 3.5 Isolation/Reunion test

#### 3.5.1 Experimental procedure

At 55, 56 and 57 days of age, piglets were confronted to an Isolation/Reunion test in order to assess their perception of the stimuli and its potential to calm the piglets after a period of stressful isolation (Fig. 1B). The test consisted in two phases. The piglet was brought individually in a trolley to the experimental room, the hatches were opened and the piglet was gently pushed into the room. It was left alone for five min, which defined the ‘Isolation’ phase. Then, one of the stimulus (‘Human’, ‘Object’) or ‘Nothing’ was shown to the piglet for 30 seconds: the door was opened with either: a) the human standing with the stool, b) the human standing with the object, or c) nothing. Finally, the second phase named ‘Reunion’ phase occurred and consisted in either a) presence of the experimenter sitting in the room on a stool and remaining immobile and quiet, b) presence of the object tied in the room, or c) isolation in the room for 270 seconds. Each piglet was thus tested three times, with one test per piglet per day. The order of the modalities (reunion with human, object or nothing) was randomized within the days and between the piglets of a same pen.

#### 3.5.2 Behavioural analyses

The tests were video recorded (camera Bosh, Box 960H-CDD) and a recorder (Noldus Mpeg recorder 2.1., The Netherlands), linked to a LCD monitor (DELL 48 model 1907FPc) which allowed us to visualize the test from an adjacent room. The location of the piglets was monitored directly during the tests and the other behaviors later from videos, both using the software The Observer XT 14.0 (Noldus^®^, The Netherlands). All behaviors used are indicated in table 1 and correspond to numbers: 1, 8-10, 14.

#### 3.5.3 Acoustic monitoring and analyses

Vocalizations produced during the Isolation/Reunion test were recorded with a C314 microphone (AKG, Austria) placed in the center of the room and at one-meter-high, connected to a MD661MK2 recorder (Marantz, USA). The vocal types were then manually annotated (grunt, squeak, bark, scream and mixed calls), after visual inspection of spectrograms on Praat^®^ software, by an expert. Only grunts were subsequently acoustically analyzed as they represented the most frequent call type that constituted a dataset of 5766 calls. A spectro-temporal analysis was performed with custom-written codes using the Seewave R package (Sueur, Aubin, and Simonis 2008) implemented in R (R Core Team 2015). After a 0.2-8 kHz bandpass filtering (‘fir’ function), a standardized grunt was detected when the amplitude crossed a 5% amplitude threshold (‘timer’ function) to measure the duration. After amplitude normalization, the following spectral parameters were calculated (‘specprop’ function, FFT with Hamming window, window length = 512, overlap = 50%): mean, median, first (Q25) and third (Q75) quartiles, interquartile range (IQR), centroid (all in Hz). The grunt dominant frequency (in kHz) was also calculated (‘dfreq’, 50% overlapping FFTs, window length = 512), which is the mean over the grunt duration of the frequencies of the highest level of energy. Parameters measuring noisiness and entropy of the grunt were: Shannon entropy (sh), Spectral Flatness (Wiener entropy, sfm) and Entropy (H) [combining both Shannon entropy and Temporal envelop entropy, length = 512, Hilbert envelop).

### 3.6 Statistical analyses

All the statistical analyses were done on R 3.3.3 (R Core Team 2015). Synthetic variables were built with Principal Component Analyses (PCA) and models were constructed to test the effect of the factors of interest. Linear or generalized mixed effect models (‘lmer’ or ‘glmer’ function, ‘lme4’ R package) were used to test two-way interactions between factors and/or continuous covariates, piglet’s identity was put as random factor (repeated measures per piglet) in all models.

#### 3.6.1 Analysis of Choice test: spatial behavior of piglets

To be able to assess and compare the behaviors during the five minutes of the Choice test and reduce the number of tested variables, a Principal Component Analysis (PCA) was done considering all behaviors directed toward each stimulus [parameters: 2-7, 11, 14 (restricted to stimulus zone) and 16, table 1] (McGregor 2005). All PCs having an Eigen value above one were kept and constituted three behavioural response scores, which cumulatively explained 81.3 % of the variability (choicePC1 – 46%, choicePC2 – 20%, choicePC3 – 16%, variable loadings, table 2). Theses scores were used as response variables for statistical analyses. The absolute values of each parameter, in several relevant conditions of the study are available in supplementary table S1. The three behavioural response scores were used as response variables in linear models testing interacting effect of the day of the test (two levels: first or second) and the stimulus (two levels Human vs. Object); the position of the human (left or right) was added as a control for choices linked to laterality (model 1). Two additional behaviors were tested as binary variables: the first approach (Human or Object, parameter 5, table 1) and whether the piglet laid down near one stimulus (presence or absence, parameter 11, table 1). To test whether the first approach depended on a stimulus, it was tested in a binomial model (Human or Object) and the effect of the day and the position of the human were put in an additive model (model 2). The number of times a piglet laid down in the proximal zone close of a stimulus was tested as a binomial variable (presence vs. absence) and a *χ*^2^ test was used.

**Table 2:** Variable loadings of the behavioural parameters used in the Principal Component Analysis in the Choice test. All Principal components (PCs) having an Eigen value above one were kept, to build behavioural response scores. The first line of the table indicates the cumulative inertia explained by the PCs. For each PC, the percentage of (left side) as well as the relative cumulative value (right side) of a given parameter is indicated. Parameters having a percentage above the uniform distribution can be considered as explanatory parameter for a given PC.

|  | Percentage per PC |  |  | Relative cumulative values |  |  |
| --- | --- | --- | --- | --- | --- | --- |
|  | choicePC1 | choicePC2 | choicePC3 | choicePC1 | choicePC2 | choicePC3 |
| Cumulative inertia | 45.8 | 65.3 | 81.3 | - | - | - |
| Number of visits in stimulus zone | 0.63 | 60.90 | 1.84 | -2.03 | 82.86 | 2.07 |
| Mean duration in stimulus zone | 24.19 | 11.56 | 1.49 | -77.59 | -15.73 | -1.68 |
| Proportion of time in stimulus zone | 23.76 | 4.38 | 0.15 | -76.21 | 5.95 | 0.17 |
| Time spent in contact with the stimulus | 22.06 | 3.22 | 0.01 | -70.77 | -4.38 | -0.01 |
| Time spent exploring when in stimulus zone | 0.22 | 17.21 | 37.35 | -0.70 | 23.41 | -41.97 |
| Latency to approach zone | 0.90 | 1.62 | 59.12 | 2.89 | -2.21 | -66.44 |
| Total time in zone | 28.25 | 1.11 | 0.03 | -90.64 | 1.51 | -0.04 |

#### 3.6.2 Analysis of Isolation/Reunion tests: spatial and vocal behavior of piglets

##### 3.6.2.1 Behavioural response scores

To be able to have comparable behaviors, between phases and stimuli, and to reduce the number of variables, behaviors were gathered and a PCA was computed (parameters: 1, 8-10, 14, table 1) (McGregor 2005). Only parameters measurable in any condition (phase of the test and type of reunion) were kept and the percentage of explained variance maximized. All PCs having an Eigen value above one were kept and constituted three behavioural response scores, which cumulatively explained 82 % of the variability (IsoReuPC1 – 32%, IsoReuPC2 – 39%, IsoReuPC3 – 11%, variable loadings, table 3). Theses scores were used as response variables for statistical analyses. The absolute values of each parameter, in relevant groups of the study are available in supplementary table S2.

**Table 3:** Variable loadings of the behavioural parameters used in the Principal Component Analysis in the Isolation/Reunion test. All Principal components (PCs) having an Eigen value above one were kept to build behavioural response score. The first line of the table indicates the cumulative inertia explained by the PCs. For each PC, the percentage of (left side) as well as the relative cumulative value (right side) of a given parameter is indicated. Parameters having a percentage above the uniform distribution can be considered as explanatory parameter for a given PC. Parameters quantifying total duration or numbers were standardised per minute.

|  | Percentage per axis |  |  | Relative cumulative values |  |  |
| --- | --- | --- | --- | --- | --- | --- |
|  | IsoReuPC1 | IsoReuPC2 | IsoReuPC3 | IsoReuPC1 | IsoReuPC2 | IsoReuPC3 |
| Cumulative inertia | 31.5 | 70.5 | 82 | - | - | - |
| Time spent standing immobile | 22.89 | 3.49 | 6.33 | -50.52 | -5.20 | -7.82 |
| Time spent looking at exit door | 2.98 | 37.10 | 13.93 | -6.57 | -55.32 | -17.21 |
| Time spent in proximal zone | 6.69 | 5.34 | 47.46 | 14.77 | 7.96 | -58.61 |
| Time spent in distal zone | 27.34 | 2.16 | 3.26 | -60.33 | 3.22 | 4.03 |
| Number of virtual zone changes | 21.90 | 5.86 | 19.76 | -48.32 | 8.74 | -24.39 |
| Time spent exploring the room | 12.00 | 25.83 | 2.60 | -26.47 | 38.52 | 3.21 |
| Latency to enter proximal zone | 6.21 | 20.22 | 6.66 | -13.69 | -30.15 | 8.22 |

##### 3.6.2.2 Acoustic scores

To be able to compare the spectro-temporal structure of grunts, two scores were built. The duration of grunts was log transformed and used as a temporal score (linear distribution). For spectral analysis, parameters previously extracted were gathered in a PCA to be able to monitor which parameters load the same way and build an acoustic score. Only one PC had an Eigen value above one, explained 83% of the variability and was named ‘Acoustic spectral score (PCac.)’. The absolute values of each parameter, in relevant groups of the study are available in supplementary table S3.

##### 3.6.2.3 Statistical models

The three behavioural response scores (IsoReuPC1, IsoReuPC2, IsoReuPC3) and the two acoustic scores (PCac. and log(duration)) were used as response variables in a linear model testing i) the two-way interaction between the type of reunion (Human/Object/Nothing) and the phase of the test (Isolation/Reunion), ii) the two-way interaction between the day of the test (1/2/3) and the phase iii) the day of the test and the type of reunion (model 3).

#### 3.6.3 Analyses of predictors for vocal expression during the reunion with the stimulus

The aim of this analysis was to search for the best predictors of vocal dynamic and grunt acoustic features, in presence of the human or the object. For this analysis, only the dataset containing the reunion phases with the Human or the Object were used, extracted from the Isolation/Reunion test.

##### 3.6.3.1 Monitoring spatial proximity toward the stimulus and time during the test

The location of the piglet in the room was divided into two categories: when the piglet was located in the proximal zone (‘Close’) and when the piglet was located elsewhere (‘Away’) to build a two level factor named ‘Location’. This parameter allowed us to track for *spatial proximity* toward the stimulus. Each period of time the piglet was Close or Away was considered as a time interval. Each time interval was numbered to track the rank of the interval along the test and the ‘Interval index’ variable was created (e.g. Close1, Away2, Close 3…).

##### 3.6.3.2 Building behavioural proximity scores toward the stimulus

Using the behavioural observations during the Choice test, *behavioural proximity scores* reflecting the closeness and exploration toward each stimulus (parameters: 2-4, 7, 12-13, 15-18, table 1) were built using two PCAs (one per type of stimulus). Only the principal component was kept in each PCA (HproxPC1 and OproxPC1) to be used as behavioural proximity score for a specific stimulus (variable loadings table 5). Only scores from day one were used, to minimize habituation effects that could occur on day two. For human, HproxPC1 explained 63% of data variability and for the object, OproxPC1 explained 47%. For further analyses, the score toward each stimulus was matched accordingly to the type of reunion the piglet was experiencing (Human vs. Object): when reunited with the human, the behavioural proximity score toward the human (HproxPC1) was used, whereas when reunited with the object, the behavioural proximity scores toward the object (OproxPC1) was used.

**Table 4:** Variable loadings of the parameters used in the Principal Component Analysis to build a spectral acoustic score. All Principal components (PCs) having an Eigen value above one were kept to build an acoustic spectral response score. The first line of the table indicates the cumulative inertia explained by the PCs. Only one PC was kept, table indicates the percentage of (left side) as well as the relative cumulative value (right side) of a given parameter. Parameters having a percentage above the uniform distribution can be considered as explanatory parameter for a given PC.

|  | Percentage on axis | Relative cumulative values |
| --- | --- | --- |
| Cumulative inertia | 83.496 | - |
| Mean | 14.820 | -98.992 |
| Centroid | 14.820 | -98.992 |
| Inter Quartile Range | 13.624 | -91.003 |
| Spectral Flatness (sfm) | 14.492 | -96.802 |
| Shannon index (sh) | 14.398 | -96.172 |
| Entropy | 13.797 | -92.159 |
| Mean Dominant frequency | 1.193 | -7.967 |
| Spectral Standard Deviation (sd) | 12.858 | -85.885 |

**Table 5:**
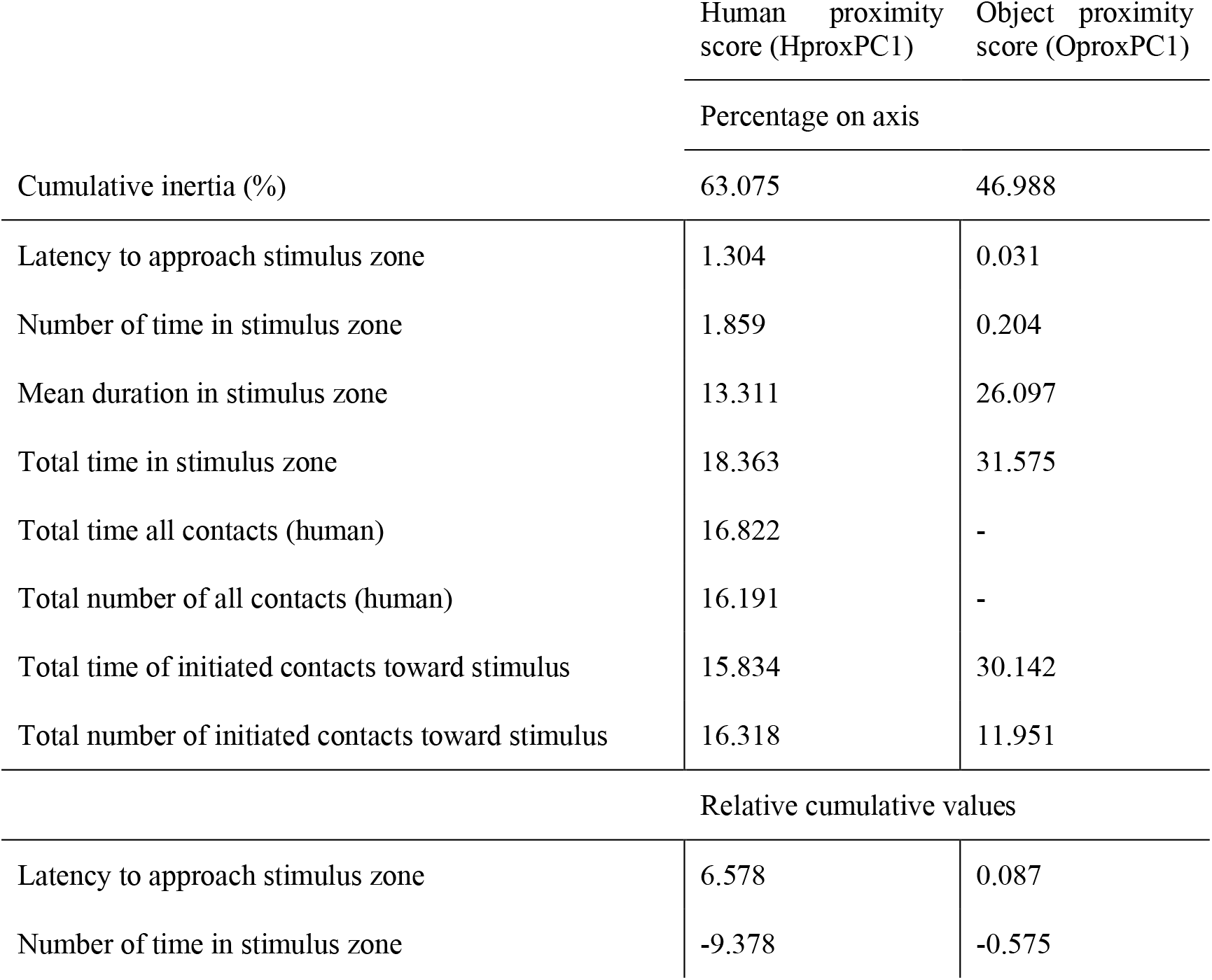
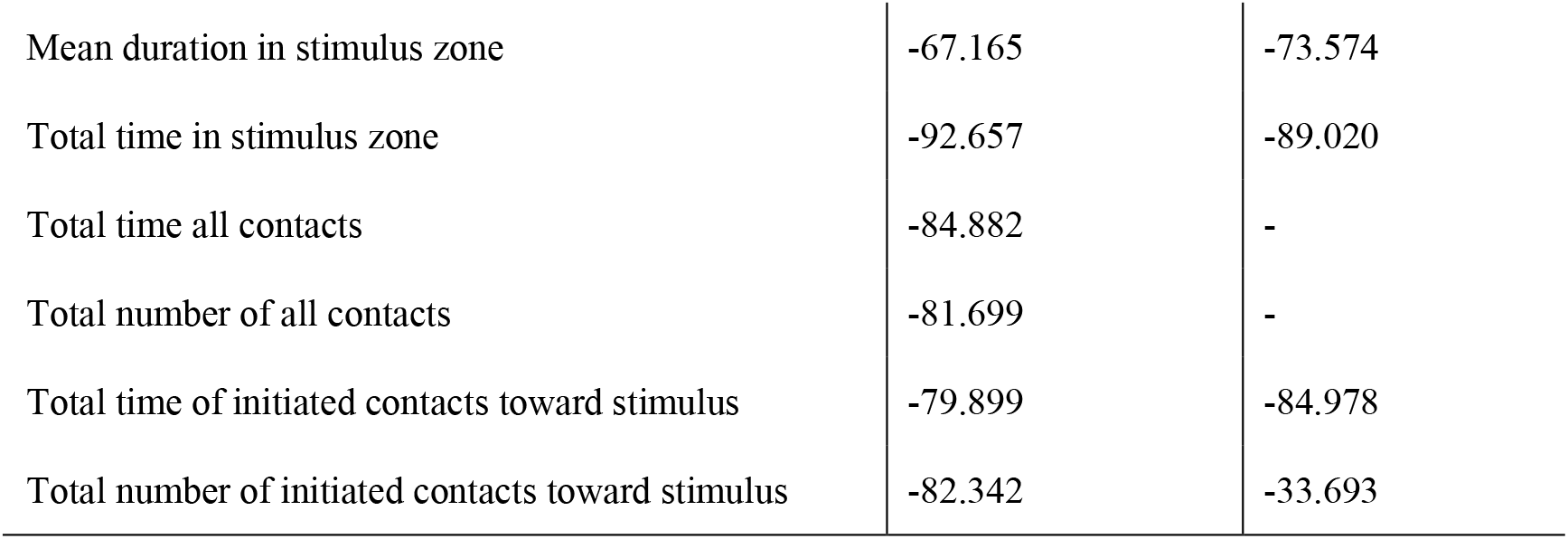
Variable loading of PCA describing piglet-stimulus behavioural proximity. Only the first Principal components was kept to create a score to be used as explanatory continuous variable. The first line of the table indicates the cumulative inertia explained by the selected PC, the percentage of (above) as well as the relative cumulative value (below) of a given parameter is indicated. Parameters having a percentage above the uniform distribution can be considered as explanatory parameter the PC. Behavioural parameters used to build these scores were extracted from the first Choice test. For statistics, the behavioural proximity score toward each stimulus was matched accordingly to the type of reunion the piglet was experiencing (Human vs. Object): when reunited with the human, the behavioural proximity score toward the human (HproxPC1) was used, whereas when reunited with the object, the behavioural proximity score toward the object (OproxPC1) was used.

|  | Human proximity<br>score (HproxPC1) | Object proximity<br>score (OproxPC1) |
| --- | --- | --- |
| Percentage on axis |  |  |
| Cumulative inertia (%) | 63.075 | 46.988 |
| Latency to approach stimulus zone | 1.304 | 0.031 |
| Number of time in stimulus zone | 1.859 | 0.204 |
| Mean duration in stimulus zone | 13.311 | 26.097 |
| Total time in stimulus zone | 18.363 | 31.575 |
| Total time all contacts (human) | 16.822 | - |
| Total number of all contacts (human) | 16.191 | - |
| Total time of initiated contacts toward stimulus | 15.834 | 30.142 |
| Total number of initiated contacts toward stimulus | 16.318 | 11.951 |
| Relative cumulative values |  |  |
| Latency to approach stimulus zone | 6.578 | 0.087 |
| Number of time in stimulus zone | -9.378 | -0.575 |
| Mean duration in stimulus zone | -67.165 | -73.574 |
| Total time in stimulus zone | -92.657 | -89.020 |
| Total time all contacts | -84.882 | - |
| Total number of all contacts | -81.699 | - |
| Total time of initiated contacts toward stimulus | -79.899 | -84.978 |
| Total number of initiated contacts toward stimulus | -82.342 | -33.693 |

##### 3.6.3.3 Model selection: searching for the best predictors of vocal expression

During the reunion with a stimulus (Human or Object), several variables could explain the vocal expression of piglets: the day of the test (3 levels), the time along the test (index of the time interval in a zone as a continuous variable), the spatial proximity of the piglet toward the stimulus (two levels: close to or away from the stimulus), the behavioural proximity of the piglet toward the stimulus (continuous behavioural proximity score) or interactions between the type of stimulus and the location, between the type of stimulus and the behavioural proximity toward the stimulus or between the type of stimulus and the time along the test. To search of the best predictors of vocal expression, five acoustic variables were used as response variables. Three variables were linear: the acoustic spectral score PCac., the duration of grunt (log(grunt duration)) and the grunt rate (number of grunt per second, calculated when the number of grunts per interval was above three (186 intervals out of 286 interval). Two non-linear variables were used: the total number of grunts (Poisson distribution), the number of times grunts were produced in series (Binomial distribution), i.e when more than one grunt was produced in a given interval. Indeed, since we used only interval containing at least three grunts to calculate the grunt rate, we needed to control we were not missing information on interval containing fewer grunts, so we used the occurrence of one grunt interval to counteract the effect of interval selection.

A full model, describing the experimental design, was built as follow (‘lmer’ or ‘glmer’ function of ‘lme4’ R package): Model 4 = Response variable ~ day + stimulus + location + Z interval index + Z behavioural proximity score + stimulus*location + stimulus*Z behavioural proximity score + stimulus*Z interval index + location*Z behavioural proximity score +(1|individual). To increase interpretability, all continuous variables (interval index and behavioural proximity scores) were scaled, so the Z score is presented every time (Schielzeth 2010), the individual was put as random factor to take into account multiple tests of the same piglet. On this full model, a model selection was performed with the ‘dredge’ function of the ‘MuMIn’ R package (Bartoń 2016), which compares all possible models built using subsets of the initial explanatory variables of the full model, including null model. Models were compared using Akaike Information Criteria corrected for small sample size (AICc). Significant models were selected when delta AICc was below two (Burnham and Anderson 2002), the weight of remaining explanatory variables was evaluated by calculating the presence or absence of the term in the remaining models (‘importance’ function). It has to be noticed that for the occurrence of one grunt intervals (Binomial model) no significant models were selected since the null model was contained in the best selected models (AICc<2). However, although it is not mentioned in the results section, the model selection table is available (supplementary material table S7).

#### 3.6.4 Tests and validation of all models and model selection

All linear models were validated by visual inspection of the symmetrical and normal distribution of the residuals (‘plotresid’ in ‘RVAideMemoire’ R package (Hervé 2016)). For generalized models, overdispersion was tested using the ‘overdisp.glmer’ function (‘RVAideMemoire’ R package, when overdispersed, a correction with the line number as random factor was used.

Anovas were computed on models to test for significant effects of explanatory variables (‘car’ R package), effect were considered significant when the p value was below 0.05. Model estimates and pairwise post hoc tests were computed using Tukey correction for multiple testing (‘lsmeans’ R package (Lenth 2016), (models 1-3)). A complete report of statistics is available as supplementary material (supplementary tables S4-S6).

For the model selection (model 4), the analysis does not give p values but rather a subset of significant models and weight of predictors. A model averaging step (‘model.avg’ function) gives the estimates of each of the predictors. The best predictors were the ones with a weight of one, meaning they were consistently present in all selected models. A complete report of all best equivalent best models is available in supplementary material (supplementary tables S7).

## 4 Results

### 4.1 Choice test between Human and Object

The PCA allowed us to extract three behavioural response scores, respectively choicePC1, choicePC2 and choicePC3 that explained cumulatively 81% of data variability (table 2).

The first behavioural response score (choicePC1, 45.8%) was negatively correlated with the mean duration in stimulus zone, proportion of time spent in stimulus zone, time spent in contact with the stimulus and the total time spent in zone. Statistics revealed a significant effect of the interaction between the type of stimulus and the day of the test (*χ*_21_ = 6.3, p = 0.012), but post hoc tests did not show any difference between groups (pairwise tests with Tukey correction, t.ratio <|2.2|, p>0.15, fig. 2A). The second behavioural response score (choicePC2, 19.5%) was positively correlated with the number of visit in stimulus zone. Statistics showed no interaction between the type of stimulus and the day of the test (*χ*_21_ = 0.7, p = 0.4), a trend for the effect of the day (*χ*_21_ = 3.3, p = 0.07) and a main effect the type of stimulus: PC2 was higher when considering the human zone compared to the object zone (*χ*_21_ = 7.3, p = 0.007, fig. 2B). The third behavioural response score (choicePC3, 16%) was negatively correlated with the time the piglet spent exploring the stimulus zone and the latency to approach the stimulus zone. Statistics showed no effect of explanatory variables on choicePC3 (Stimulus: *χ*_21_ = 1.5, p = 0.2, Day: *χ*_21_ = 0.6, p = 0.5). We counted the number of times the human zone or the object zone was first approached by the piglet during the test. Statistic showed a trends for the effect of the day of the test: piglets tended to approach first the object zone more often the second day than the first day of the test (*χ*_21_ = 3.4, p = 0.06, fig. 2C). Last, we counted the number of times piglets laid down near a stimulus zone, a *χ*^2^ showed a significantly different distribution of occurrences of this behavior, that only occurred in the human zone (by nine individuals out of 24) and none in the object zone (*χ*^2^_1_= 12.8, p < 0.001, fig. 2D). The position of the human in the room (left or right side) was included in all models and never showed a significant effect (see supplementary table S4 for full report).

**Figure 2:**
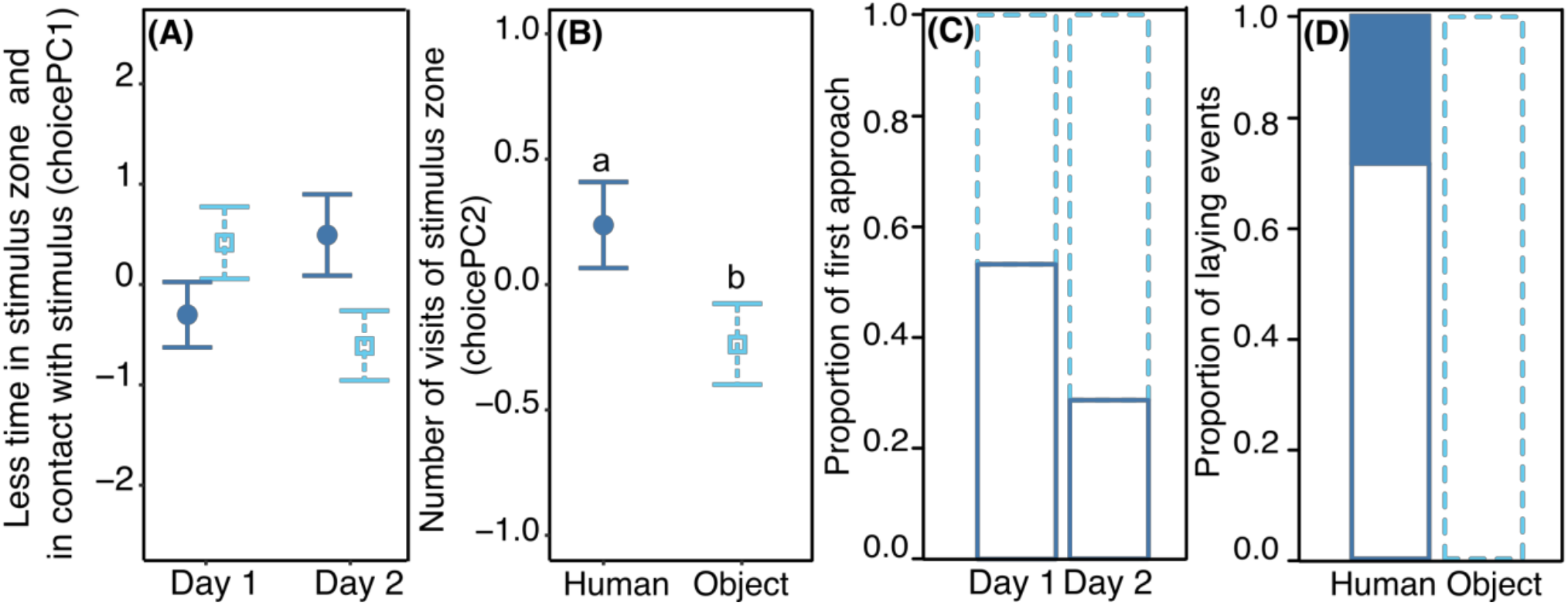
Behavioural response in the Choice test. Mean (± se) of the two behavioural response scores from the PCA analysis, choicePC1 (A) and choicePC2 (B), toward the two possible stimuli: either Human (full dark blue circles) or Object (empty light blue squares). A: Significant interaction between the stimulus and the day of the test but no differences revealed between groups in post hoc tests. B: significant effect of the stimulus on choicePC2. C: first approach of the piglet toward one of the stimulus either human (solid dark blue bars) or object (dotted light blue bars), indicated as proportion of the 24 piglets tested twice (day 1 and day 2). D: Proportion of times piglets that laid down during the test either in the human zone or in the object zone. Different letters indicated significantly different groups (p<0.05). All model estimates, anova tables and results of post hoc tests are available in supplementary tables S4-S6.

### 4.2 Isolation/Reunion test

#### 4.2.1 Variation of piglets’ behavior when they are reunited with a human or an object

For the Isolation/Reunion test, a PCA allowed us to extract three behavioural response scores, respectively IsoReuPC1, IsoReuPC2 and IsoReuPC3 that explained cumulatively 82% of data variability (table 3).

The first behavioural response score (IsoReuPC1, 31.5%) was negatively correlated with the time spent immobile, the time spent in distal zone and the number of changes of virtual zone. Statistics revealed a significant interaction between the type of reunion and the phase of the test (*χ*_22_ = 16.6, p < 0.001, fig. 3A). During the isolation phase, no significant difference was found between groups (pairwise comparisons human/object/nothing, |t.ratio| < 0.7, p > 0.9) whereas during the reunion phase, the three type of reunions differed significantly in PC1 values (human vs. object: t.ratio = 3.1, p = 0.03, human vs. nothing: t.ratio = 6.3, p < 0.001, object vs. nothing: t.ratio = 3.3, p = 0.02). The reaction to each reunion type did not have the same amplitude too. When piglets were not reunited with a stimulus, statistics did not show differences between the isolation and the reunion phases (isolation vs reunion with nothing: t.ratio = 0.6, p > 0.9), whereas when reunited with a stimulus, IsoReuPC1 showed a significant increase that was stronger with the human (isolation vs. reunion, t.ratio = −6.3, p < 0.001) than with the object (isolation vs. reunion, t.ratio = −3.2, p < 0.03).

**Figure 3:**
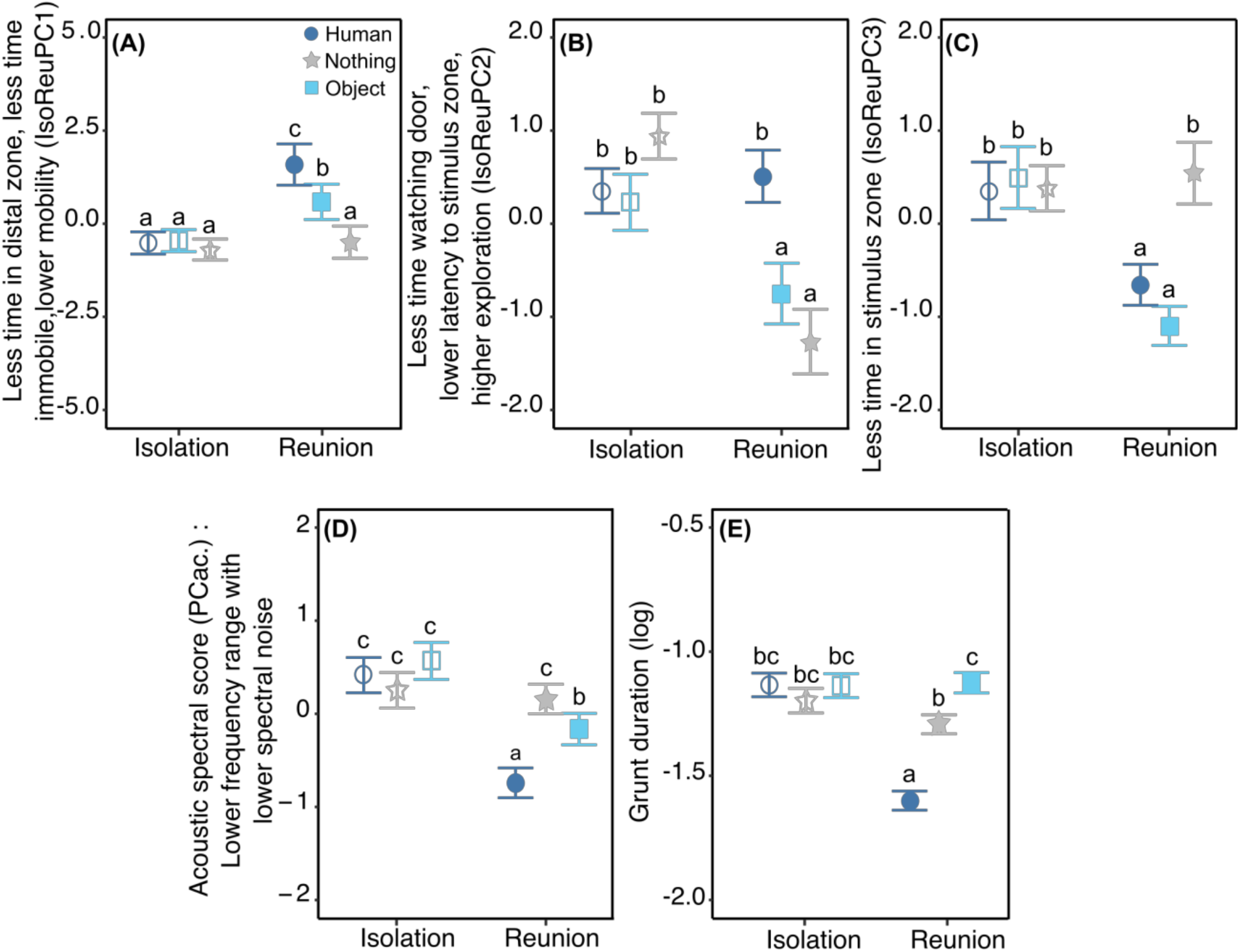
Behavioural (A to C) and vocal (D, E) responses to the Isolation/Reunion test. Mean (± se) of behavioural and acoustic scores, according to the stimulus (Human = dark blue circles, Object = light blue squares, Nothing = grey stars), the phase of the test (Isolation = empty symbols or Reunion = full symbols) and/or the day of the test (day 1, 2 or 3). A-C: Significant interaction between the type of reunion and the phase of the test for the three behavioural response scores IsoReuPC1, IsoReuPC2, IsoReuPC3, respectively. D-E: Significant interaction between the type of reunion and the phase of the test on the two acoustic score: the acoustic spectral score (D) and the logarithm of grunt duration (E). Different letters show significantly different groups (p < 0.05). All model estimates, anova tables and results of post hoc tests are available in supplementary tables S4-S6.

The second behavioural response score (IsoReuPC2, 39%) was positively correlated with the time spent exploring the room and negatively correlated with the time spent looking at the entrance door and the latency to enter the proximal zone of the stimulus. A significant interaction was found between the type of reunion and the phase of the test (*χ*_22_ = 41. 5, p < 0.0001, fig. 3B). During the isolation phase, no significant difference was found between groups (pairwise comparison human/object/nothing, |t.ratio| < 2.7, p > 0.08) whereas during the reunion phase, the two types of stimuli differed significantly (human vs. object: t.ratio = 4.9, p < 0.001), as well as the reunion with the human compared to nothing (t.ratio = 6.8, p < 0.001), but no difference was found when comparing reunions with the object or nothing (t.ratio = 2.0, p = 0.37). The reaction to the three types of reunions differed too: from isolation to reunion phase, no difference was found in IsoReuPC2 when piglets were reunited with the human (t.ratio = −0.6, p = 0.9), whereas PC2 decreased significantly when piglets were reunited with the object or with nothing (object: t.ratio = 3.8, p = 0.003, nothing: t.ratio = 8.5, p < 0.001).

The third behavioural response score (IsoReuPC3, 11.5%) was negatively correlated with the time spent in the stimulus zone. Statistics showed a significant interaction between the type of reunion and the phase of the test on IsoReuPC3 (*χ*_22_ = 36.4, p < 0.001, fig. 3C). During the isolation phase, no significant difference was found between groups (pairwise comparison human/object/nothing, |t.ratio|<0.7, p>0.9). During the reunion phase, for piglets being reunited with nothing, IsoReuPC3 differed significantly compared to being reunited with a stimulus (human vs. nothing: t.ratio = −5.7, p < 0.001, object vs. nothing: t.ratio = −7.8, p < 0.001) but IsoReuPC3 between the two types of stimuli did not differ (human vs. object: t.ratio = 2.1, p = 0.3). The reaction to the three types of reunions differed too: from isolation to reunion phase, no difference was found in IsoReuPC3 when piglets were not reunited with a stimulus (reunion with nothing: t.ratio = −0.8, p = 0.9), whereas IsoReuPC3 decreased significantly when piglets were reunited with the object or with the human (object: t.ratio = 7.6, p < 0.001, human: t.ratio = 4.8, p < 0.001).

The day of the test did not show any effect on IsoReuPC2 and IsoReuPC3 (*χ*_22_ = 0.9, p = 0.6, *χ*_22_ = 0.2, p = 0.9 respectively) but was significantly higher for IsoReuPC1 from day one to day three (*χ*_22_ = 10.1, p = 0.007, supplementary figure S1A). Post hoc test showed the differences in IsoReuPC1 were progressive along days (pairwise comparison, day 1 vs. day2: t.ratio = −2.4, p = 0.05, day 1 vs. day 3: t.ratio =-3.0, p = 0.009, day 2 vs. day3: t.ratio = −0.6, p = 0.79).

#### 4.2.2 Variation of piglets’ grunts acoustic features when they are reunited with a human or an object

All 5766 grunts produced during the test were analyzed using two acoustic scores: the logarithm of grunt duration and a spectral score, computed from the PCA of frequency and noise parameters of the calls (acoustic spectral score PCac., variable loading table 4): the greater the score, the lower the frequency and the lower the spectral noise in the grunt.

Concerning the spectral acoustic score (PCac.), a significant interaction was found between the type of reunion and the phase of the test (*χ*_21_ = 45.1, p < 0.001, fig. 3D). During the isolation phase, no difference was found between groups (pairwise comparison during isolation, human/object/nothing: |t.ratio| < 1.9, p > 0.4) whereas during the reunion phase significant differences were found between groups (pairwise comparisons during reunion, human vs. object: t.ratio = −4.9, p < 0.001, human vs. nothing: t.ratio = −9.2, p < 0.001, nothing vs. object: t.ratio = 3.7, p = 0.003). The reaction to each of the reunion types did not have the same amplitude too. When piglets were subjected to another isolation, statistics did not show differences between the isolation and the reunion phase (t.ratio = 0.03, p = 1), whereas when reunited with a stimulus, PCac. showed a significant decreased that was stronger with the human (t.ratio = 9.3, p < 0.001) than with the object (t.ratio = 5.3, p < 0.001). Statistics also showed a significant interaction between the type of reunion and the day of the test (*χ*_21_ = 26.8, p < 0.001) but post hoc tests revealed no significant pairwise comparisons (|t.ratio| < 1.6, p > 0.8, see supplementary table S4-S6 and supplementary fig. S1B).

Grunt duration showed a significant interaction between the type of reunion and the phase of the test (*χ*_22_ = 210.1, p < 0.001, fig. 3E). During the isolation phase, no difference was found between groups (pairwise comparison during isolation, human/object/nothing: |t.ratio| < 2.6, p > 0.09) whereas during the reunion phase significant differences were found between groups (pairwise comparisons during reunion, human vs. object: t.ratio = −19.5, p < 0.001, human vs. nothing: t.ratio = −16.7, p < 0.001, nothing vs. object: t.ratio = −3.9, p = 0.003). The reaction to each of the reunion types differed too. When piglets were subjected to another isolation or reunited with the object, statistics did not show differences between the isolation and the reunion phase (pairwise comparisons isolation vs. reunion object/nothing: |t.ratio| < 0.6, p > 0.6), whereas when reunited with the human, grunt duration decreased significantly (pairwise comparison isolation vs. reunion, human: t.ratio = 9.3, p < 0.001). Last, statistics also revealed a significant main effect of the day of the test (*χ*_22_ = 20.0, p < 0.001): grunt duration decreased as the day of the test increased, especially between the first two days (pairwise comparisons, day 1 vs. day2: t.ratio = 3.9, p < 0.001, day 1 vs. day 3: t.ratio = 2.6, p = 0.03, day 2 vs. day 3: t.ratio = −1.2, p = 0.4, supplementary figure fig. S1C).

### 4.3 Effect of proximity to stimulus on vocal expression

The four following acoustic variables: total number of grunts, grunt rate, duration of grunts (log(grunt duration)) and spectral acoustic score (PCac.) may be predicted by the context (the type of stimulus), the spatial proximity to the stimulus (location in the room), variables independent from the stimuli (day, time along the test, described by the interval index) or the experience piglets previously had with the stimuli. To quantify the experience piglets had with each stimulus (closeness and exploring), a behavioural proximity score resulting two from principal component analyses were built (table 5) and one was selected per type of reunion: ‘behavioural proximity score’ corresponded to the opposite of HproxPC1/OproxPC1 (respectively for reunion with the human or the object) and was positively correlated with the time spent in contact with and near the stimulus. After model comparison and selection of the best equivalent models, the weight of predictors as well as the estimates of the averaged resulting model were calculated (tables 6 and 7, respectively, full selected models in supplementary table S7).

**Table 6:** Weight of predictors for each response variable. The number of equivalent best models (N), and each term of the full initial model are indicated in columns: ‘Stim’ for stimulus type, Day, ‘Loc’ for location in the room (Close to or Away to the stimulus), ‘Int. Index’ for interval index, ‘Behav Prox’ for behavioural proximity score, as well as relevant interactions between explanatory variables. Only weights different from zero are indicated. For the total number of grunts, since the variable is a sum of all intervals per day, location, individuals and type of stimulus, the interval index was not included in the full model and referred as ‘NA’. The best predictors are the one consistently appearing in all equivalent selected models so the ones having a weight of ‘1’.

|  | N | Stim | Day | Loc | -Behav<br>Prox | Stim<br>*Day | Stim<br>*Loc | Stim<br>*-<br>Behav<br>Prox | Loc *-<br>Behav<br>Prox | Int<br>Index | Stim<br>*Int<br>index |
| --- | --- | --- | --- | --- | --- | --- | --- | --- | --- | --- | --- |
| Total Number of grunt-Poisson | 2 | 1.00 | 1.00 | 1.00 | 1.00 | - | 1.00 | 1.00 | 0.47 | NA | NA |
| Grunt rate (log) | 2 | 1.00 | - | - | 0.30 | - | - | - | - | - | - |
| Grunt duration (log) | 6 | 1.00 | 1.00 | 1.00 | 1.00 | - | 1.00 | 0.24 | 0.25 | 1.00 | 0.51 |
| Acoustic spectral score<br>(PC1ac.) | 9 | 1.00 | 1.00 | 1.00 | 1.00 | 0.36 | 1.00 | 0.44 | 0.13 | 1.00 | 0.24 |

**Table 7:**
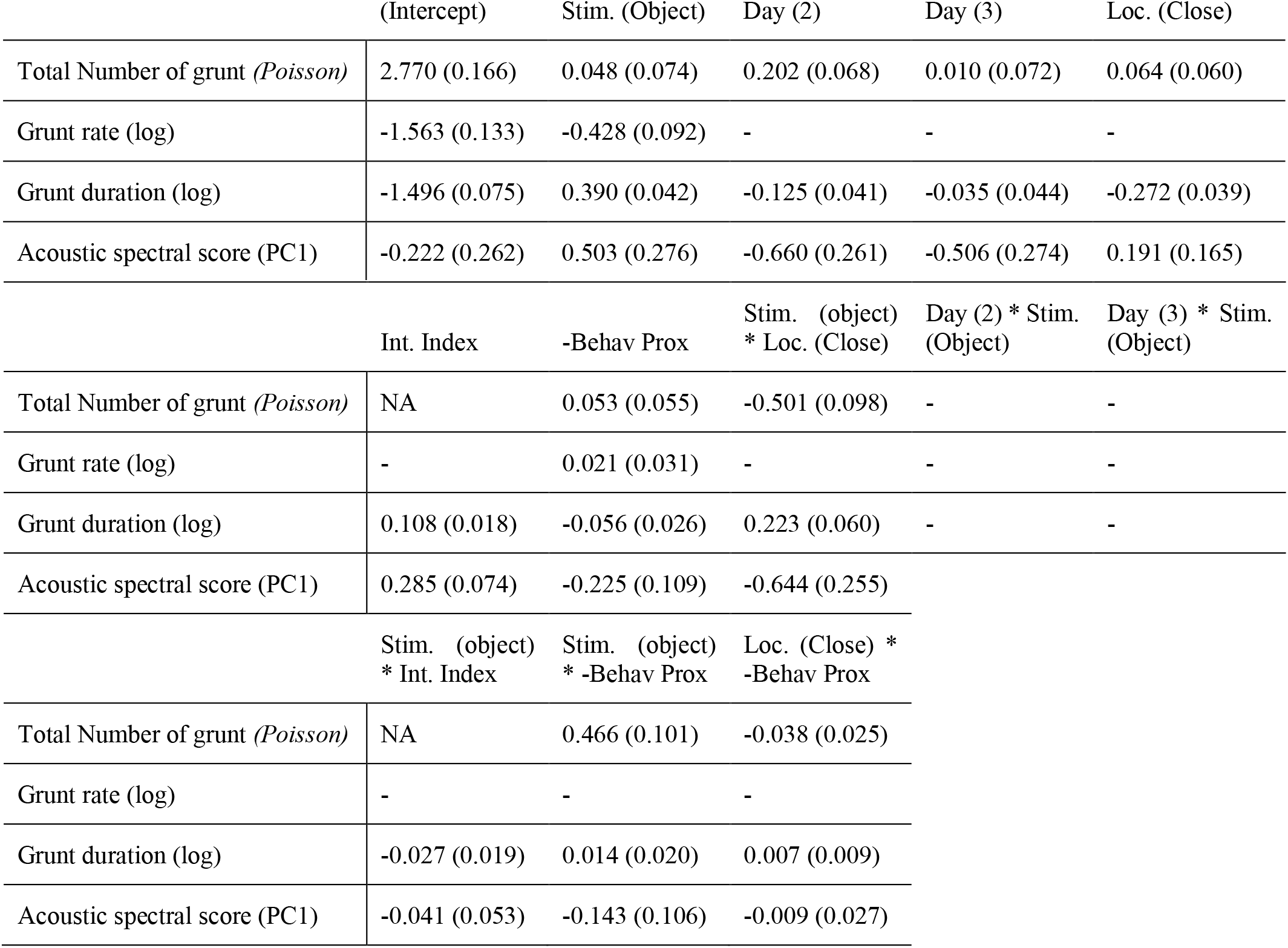
Estimates (standard error) of terms contained in the equivalent best selected models. When the term is a factor, the estimate is indicated for one level and the absolute value has to be calculated using the estimate of the intercept, the level compared is indicated. When the term is a continuous covariate, the estimate of the slope is indicated, notice that to increase interpretability, all continuous were scaled so the value is for the Z score. ‘-’ indicates the term was not selected in the best equivalent model. ‘NA’ indicates the term was not in the full model prior to the model selection.

The model selection showed the total number of grunts was predicted by the interactions between the type of stimulus and the location of the piglet in the room, as well as the interaction between the type of stimulus and the behavioural proximity score (table 6). Thus, a lower number of grunts was likely to predict the piglet was reunited with the object, and spatially close to it (fig. 4B). In addition, when reunited with the object, the higher the behavioural proximity score (-OproxPC1), the higher the probability of producing more grunts (fig. 4A), but not with the human. Concerning grunt rate, the type of stimulus was the only consistent predictor (table 6): when piglets were reunited with the human, the rate of grunt was higher than when reunited with the object (fig. 4C).

**Figure 4:**
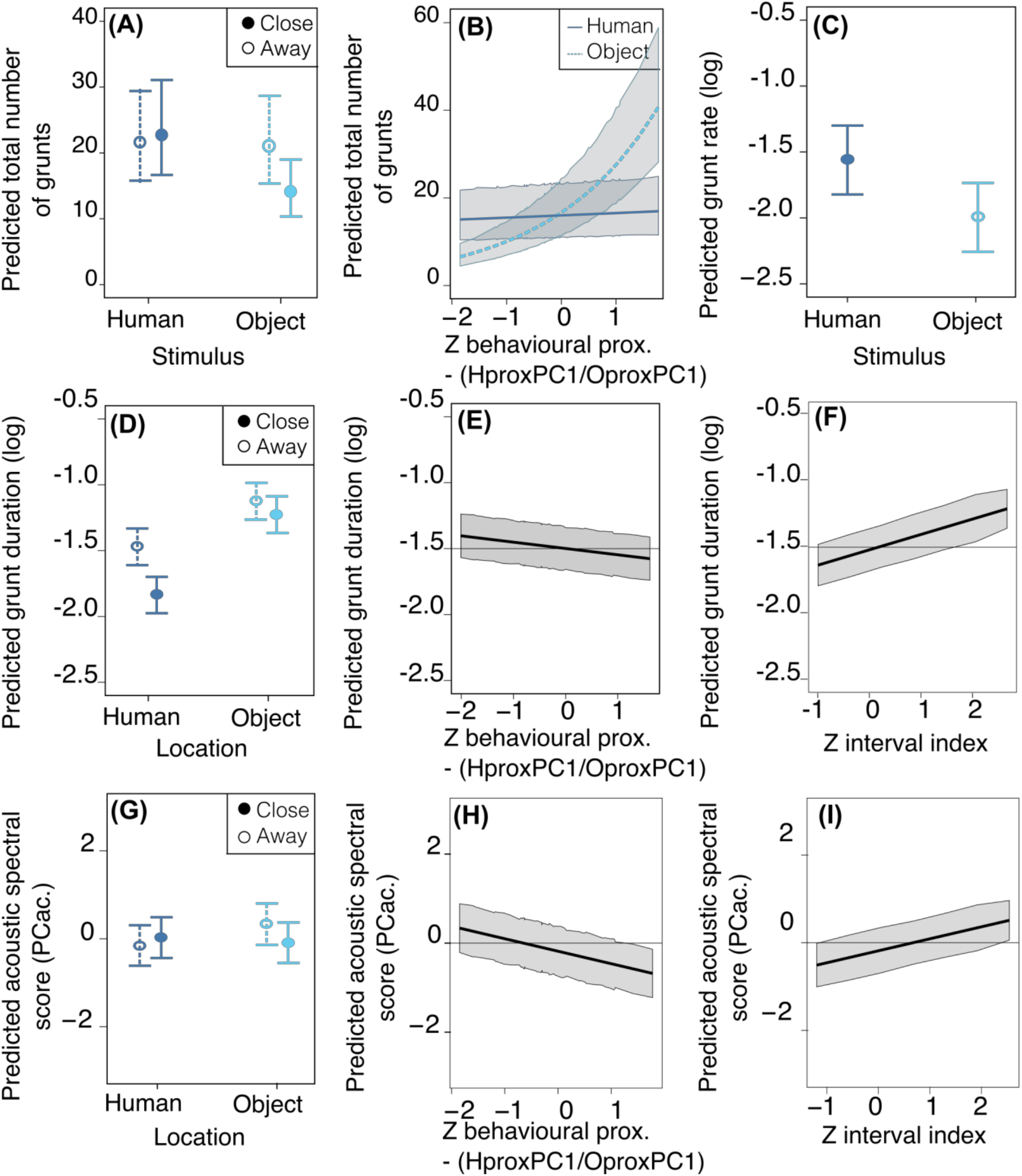
Mean estimates and 95% confidence interval of best predictors of vocal expression: number of grunts (A,B), grunt rate (C), grunt duration (D,E,F) and acoustic spectral score PCac. (G, H, I), depending on stimulus (cyan for object, dark blue for human, location in the room, behavioural proximity and time along the test. Best predictors are represented for illustrating the range and directions of effects. Location (A,D,G): whether the piglet was located close to the stimulus (solid lines) or away from it (dotted lines). Behavioural proximity score (B, E, H) was scaled for the statistical analysis so the Z score is represented (see composition of scores in table 5). (A-D, G) Type of stimulus: whether the reunion was with the human (dark blue solid circles and lines) or the object (light blue empty circles and dotted lines. Time along the test (F, I) is quantify with the interval index along the test and is scaled, so the Z score is represented. Plots were generated using the averaged best model resulting from the model selection (models having delta AICc below 2 and predictors having a weight of 1), for which the estimates (se) are in table 7, the full selection model table is available in supplementary table S7.

Considering the acoustic structure of grunts (duration and spectral acoustic score PCac.), both descriptors were best predicted by the interaction between the location in the room and the type of stimulus, the behavioural proximity score, the interval index and the day (table 6). The probability of having shorter grunts was higher when reunited with the human and close to her (fig. 4D). In addition, the higher the behavioural proximity score, the higher the probability of having shorter grunts (fig. 4E). The probability of having longer grunts increased as the time of the test increased (interval index, fig. 4F) with no interaction with the type of stimulus or location. Last, as the day of the test increased, the probability of having shorter grunt increased (slope estimate ± se: −0.13 ± 0.04 and −0.04 ± 0.04, respectively for day two and three, table 7), with no interaction with the type of stimulus. Concerning the acoustic spectral score, (fig. 4E, table 7): the probability of producing grunts with a lower acoustic spectral score depended of the type of stimulus and the spatial proximity, as the acoustic spectral score was more likely to decrease when approaching the object but not the human (fig. 4G).The higher the behavioural proximity score, the higher the probability of producing grunt with a lower acoustic spectral score, independently from the type of stimulus and location in the room (fig. 4H, table 6). As the time during the test increased, the probability of producing grunts with a higher acoustic spectral score increased, independently from the type of stimulus or location (fig 4I). Last, as the day of the test increased, the probability of producing grunts with a lower acoustic spectral score increased independently for the type of stimulus (slope estimate ± se: −067 ± 0.26 and −0.51 ± 0.27, respectively for day two and three, table 7).

## 5 Discussion

### 5.1.1 No evidence of a preference toward a stimulus but specific human directed behaviours

In a V shaped apparatus Choice test, comparing the time spent close to and in contact with each of the stimulus (first behavioural response score, choicePC1) or the latency to reach the stimulus zone and exploring the stimulus zone (third behavioural response score, choicePC3), did not lead to significant differences between the types of stimuli. No evidence for a preference in the first approach could be found too. So this test did not allow to conclude for a preference for one of the stimuli. The novel conformation of the testing pen compared to what pigs used to know (open pen) may have attracted their attention more than the familiar stimuli present. Using the home pen to test for preference, like done in mice for instance (Blom et al. 1992) may have led to different results, even if the technical procedures would have been much more complicated. Indeed, male mice may show preferential attraction for different enrichment stimuli in such a situation (Loo et al. 2016). No particular negative behaviors associated to fear or stress related were recorded during the test, thus the situation itself may not have been negative for our piglets. In addition, the absolute time spent close to each of the stimulus (between 73.9s and 100s out of 300s) or in contact with the stimulus (between 28.5s and 67.8s out of 300s, supplementary table S5) were high enough to conclude that the stimuli were attractive. However in our study, two differences came out between the human and the object. The piglets entered more often the human zone than the object zone (second behavioural response score, choicePC2) and the number of times piglets laid down near the stimulus was human zone specific. Laying down is a sign of absence of stress in pigs (Goumon and Špinka 2016), and the location was not by chance here. We may hypothesise that the human had reassuring effects, as it has been found in studies on other farm animal species (Rushen et al. 2001; Tallet, Veissier, and Boivin 2005). This will need to be confirmed by monitoring hear rate and its variability.

### 5.1.2 Behavioural evidence for positive attractivity of both a manipulable object and the familiar human

Isolation/reunion tests allowed us to show differential responses according to the stimulus piglets were reunited with. Behavioural measures showed that stimuli were attractive for the piglets with a decrease in the time spent in the zone distant from the stimuli and an increase in the time spent in the stimulus zone when piglets were reunited with the human or the object compared to remaining alone in the experimental room. Also piglets remained less immobile inactive, a behavior associated with the vigilance response, and they had a lower locomotor activity when one stimulus was present. These changes in locomotor activity have to be explained along with the time spent near the stimulus and are in line with the hypothesis of attractivity to the stimuli. Beyond this general changes in behaviour, piglets expressed discriminative behaviours according to the stimulus present. Indeed, in response to a reunion with the human compared to the object, piglets were quicker to enter the stimulus zone, expressed a lower mobility and a higher exploration time. In response to a reunion with the object, piglets spent more time watching the exit door than exploring the room, a response equivalent to the reunion with nothing (i.e. isolation). As a consequence, results may show the presence of the familiar human may prevent the piglets from expressing stress responses (more vigilance behaviour and less exploration), a hypothesis strengthened by the fact that being reunited with the object or nothing seems equivalent in terms of vigilance behviors.

### 5.1.3 Acoustic evidence of a high arousal positive emotional state with the human and a low arousal emotional state with the object

We predicted that if vocalisations allow expression of the emotional state of the piglets, acoustic scores should be different when piglets would be reunited with a stimulus compared to nothing. In reaction to the reunion with the familiar human, the duration of grunts decreased and this was not the case with the object or when piglets remained alone. Shorter vocalisations have been associated with positive contexts compared to negative ones in many species (Briefer 2012), and specially shorter grunts in pigs (Friel et al. 2019; Briefer et al. 2019). We can compare the absolute values of grunt duration from the present study (250 ± 180ms with human, 380 ± 220ms with object, 330 ± 210ms isolated, supplementary table S3) and other studies (negative vs. positive context (Briefer et al. 2019): 480 ms vs. 280 ms; negative vs. positive context (Friel et al. 2019): ~430ms vs. ~350ms; anticipation of social reunions with pen mates (Villain et al. 2020): ~240ms. Although comparisons has to be taken precociously due to material and methodological specificity of each study, the range of values we obtained with the human here are in the range of positively perceived situations. Behavior and acoustic together may allow to conclude that being reunited with the human leads to a more positive context than with the object. Since the human has been associated with positive tactile contacts, known to promote a positive state (Tallet et al. 2014; 2018), approaching the human may engage piglets in a positive anticipation of positive tactile interactions. The object rather leads to an emotional state having a valence comparable to isolation, even if it is attractive to some extent. During reunion with either the object or the human, the spectral acoustic score of piglets grunts decreased: grunts were composed of higher frequencies and a higher noise component and this effect was greater with the human compared to the object. Changes in spectral components in response to changing contexts may be associated with the arousal of situations in mammals (Ehret 2018; Villain et al. 2020). This may underline that the reunion with a stimulus promotes emotional states of high intensity in piglets, especially the reunion with a human. Villain et al. (2020) showed piglets were able to rapidly change the spectral properties of their grunts when anticipating positive events. The anticipation of a reunion with familiar conspecifics led to noisier grunts whereas the anticipation of a reunion with a familiar human associated with positive contacts, led to higher pitched grunts. In the present study, frequency and noise components of the grunt are closely intercorrelated, so it is not possible to discriminate between the two. To summarize, for isolated piglets, being reunited with a familiar human induces a high arousal positive emotional state, while being reunited with a familiar object induces a low arousal emotional state. Thus, human positive interaction seems to be more valuable as an enrichment for piglets. This may result from the relationship created through the numerous sessions of positive vocal and tactile interactions, as already shown in previous studies (Tallet et al. 2014; Brajon, Laforest, Schmitt, et al. 2015). An inanimate object may not acquire similar properties. As a consequence promoting social or pseudo-social enrichment in pigs is a good way to enhance their welfare.

### 5.1.4 Experience and spatial location predict differences in spectro-temporal features of grunts depending on the stimulus

To go further we studied which variables predicted vocal production. From the model selection, we found that the type of stimulus (object or human) was among the best predictors of vocal expression (number of grunts, grunt rate, duration, and spectral score) and was the only consistent predictor explaining the temporal dynamic (grunt rate). Being reunited with a human (and not an object) is associated with more vocal production and at a higher rate. Morton (1977) explained that the rhythm of a behavior can be positively linked to motivation of the producer, and thus a higher arousal. Villain et al. (2020) showed piglets had a higher grunt rate when anticipating the arrival of conspecifics, compared to a familiar human. Here, we would interpret the result in the direction of a higher motivation toward the human compared to the object.

Being reunited with the human and close to them is likely to induce shorter grunts whereas being reunited with the object and close to it is likely to induce a fewer number of higher frequency and noisier grunts. This is in line with the positive state induced by the human compared to the object.

The behavioural proximity, behavioural score associated with the number of interactions and the time spent in contact with or near the stimulus was a consistent predictor for both acoustic scores. Independently from the type of stimulus, the higher the behavioural proximity to the stimulus was, the higher the probability for producing shorter grunts with higher frequencies and noise components was. This raises the question of the possibility to monitor the degree of behavioural proximity to an enrichment using the structure of grunts (Briefer et al. 2019).

The time during the test was also a predictor of the spectro temporal features: the later in the test, the higher the probability of producing longer, lower pitched and less noisy grunts was (effect of interval index). Positive effect of stimulus presence may be attenuated with time, and/or negative effects of isolation from penmates may increase. This was independent from the type of stimulus, we can hypothesize that during the test, since the human do not interact with the piglet (as it would outside the test) make the human closer to an object and piglets may habituate to the stimulus. It would be interesting to investigate whether interacting with the piglet during an Isolation/Reunion test may prolong the positive effect of the reunion with a familiar human after a five minute isolation. Last, along days, grunts were more likely to be shorter, higher pitched and noisier, independently from the type of stimulus. This may have to be linked to habituation to the protocol along test.

### 5.1.5 A familiar human as enrichment: implication for pig welfare

Reunion with a familiar object or human led to an attraction toward the stimulus and repeated contacts, as well as a decrease in vigilance behavior. These parameters are in line with the definition of what a ‘environmental enrichment’ should promote (Godyń, Nowicki, and Herbut 2019; Newberry 1995). Both stimuli we provided can be considered as enrichments. Being with a human and/or close to the human provokes higher degrees of behavioural changes in piglets (both spatial and vocal) and specific behavioural postures (laying), associated with positive states. Regarding vocal behavior, although we showed that the behavioural proximity to the stimulus and vocal responses correlated, only the human presence led to positive shorter grunts during the reunion, and not the object. Regarding enrichments, it has been pointed out that the novelty is a paramount feature to promote a long term positive context and delay habituation effects (Trickett, Guy, and Edwards 2009). It is possible that interactions with the human may allow this feature, indeed, a human is moving, talking and is not likely to reproduce exactly the same gesture, at the same rhythm, which may participate in keeping a higher level of stimulation than an object may do. More studies are needed to better describe what are the human most efficient signals and behaviors that promote positive emotional states using a multimodal approach: voice? (Bensoussan et al, 2020), shape? Facial expression? (e.g. goats (Nawroth et al. 2018):), facial cues (Wondrak et al. 2018) or odours? (review: (Nielsen 2018), combinations of factors? (Tanida and Nagano 1998).

To conclude, using behavioural and vocal monitoring, this study showed that a manipulable object and a familiar human can be attractive for weaned piglets out of their rearing environment. A familiar human may enhance a positive and high arousal emotional state when the piglet is alone, mimicking reassuring effects. Humans may be considered as enrichment in the piglets’ environment and more studies should consider pseudo social interactions between humans and piglets to enhance welfare. In order to be applicable on a larger scale, the kinetic of human-piglet relationships needs to be better understood, as well as the most efficient signals triggering positive emotional states in piglets.

## Supporting information

Supplementary data all

## 7 Author Contributions

Conceived and designed the experiment (A.V., C.T., M.L.). Performed the experiment (A.V., C.G., M.L.). Collection and edition of the acoustic and behavioural data (A.V., M.L., C.G.). Statistical analyses (A.V.). Contributed to the writing of the manuscript (A.V., C.T., M.L.).

## 9 Data Availability Statement

Data sets have been deposited to the online repository of INRAE: Villain, Avelyne S., Lanthony, Mathilde, Guérin, Carole, Tallet, Céline, 2020, “Manipulable object and human contact: preferences and modulation of emotional states in weaned piglets”, https://doi.org/10.15454/GDLDBH, Portail Data INRAE (Villain et al. 2020). They will be accessible once the article is published.

## 10 Acknowledgments

We acknowledge all the technical staff at UEPR: especially Patrick Touanel, who largely participated in handling the piglets. We thank Eric Siroux who helped building the acoustic chamber at the beginning of the experiment and Remi Resmond for great discussion about statistics.

