## Supplementary data all for "Manipulable object and human contact: preferences and modulation of emotional states in weaned piglets"

### 12 Supplementary tables

**Supplementary table S7**: Absolute values per group of parameters loaded to the Principal component analysis for the choice test. For each parameter, the number of observations (N) and the group –defined by a stimulus (Human or Object) during a day (Day 1 or Day2), is indicated. For each parameter, the mean, standard deviation (sd) and 95% confidence interval (ci) are indicated

| | | | **Number of visits in zone** | | | **Mean duration in zone (s)** | | | **Proportion of time in zone** | | | **Time spent in contact to the stimulus (s)** | | |
|---|---|---|---|---|---|---|---|---|---|---|---|---|---|---|
| **Stim** | **Day of test** | **N** | **mean** | **sd** | **ci** | **mean** | **sd** | **ci** | **mean** | **sd** | **ci** | **mean** | **sd** | **ci** |
| Human | Day 1 | 24 | 5.71 | 1.99 | 0.84 | 19.04 | 11.28 | 4.76 | 0.57 | 0.21 | 0.09 | 44.50 | 29.98 | 12.66 |
| Human | Day 2 | 24 | 4.33 | 1.86 | 0.78 | 20.12 | 15.29 | 6.45 | 0.47 | 0.22 | 0.09 | 28.48 | 26.86 | 11.34 |
| Object | Day 1 | 24 | 4.50 | 1.56 | 0.66 | 17.46 | 12.45 | 5.26 | 0.43 | 0.21 | 0.09 | 49.57 | 46.77 | 19.75 |
| Object | Day 2 | 24 | 4.04 | 1.85 | 0.78 | 29.28 | 27.92 | 11.79 | 0.53 | 0.22 | 0.09 | 67.70 | 50.53 | 21.34 |

| | | | **Total time in zone (s)** | | | **Time spent exploring in zone (s)** | | | **Latency to approach zone (s)** | | |
|---|---|---|---|---|---|---|---|---|---|---|---|
| **Stimulus** | **Day of test** | **N** | **mean** | **sd** | **ci** | **mean** | **sd** | **ci** | **mean** | **sd** | **ci** |
| Human | Day 1 | 24 | 100.05 | 53.23 | 22.48 | 9.01 | 9.35 | 3.95 | 14.61 | 8.09 | 3.42 |
| Human | Day 2 | 24 | 82.19 | 55.04 | 23.24 | 13.31 | 11.86 | 5.01 | 28.45 | 41.59 | 17.56 |
| Object | Day 1 | 24 | 73.93 | 48.56 | 20.50 | 9.56 | 10.60 | 4.48 | 20.15 | 25.42 | 10.74 |
| Object | Day 2 | 24 | 97.30 | 65.08 | 27.48 | 16.66 | 23.64 | 9.98 | 12.45 | 16.32 | 6.89 |

**Supplementary table S8**: Absolute values per group of parameters loaded to the Principal component analysis for the Isolation/Reunion test. For each parameter, the number of observations (N) and the group –defined according to a stimulus (Human or Object) and/or a day (Day 1 or Day2) and/or a phase of a test is indicated. For each parameter, the mean, standard deviation (sd) and 95% confidence interval (ci) are indicated.

| | | | Time spent standing immobile (per min.) | | | Time spent looking at exit door (per min.) | | | Time spent in proximal zone (per min.) | | | Time spent in distal zone (per min.) | | |
|---|---|---|---|---|---|---|---|---|---|---|---|---|---|---|
| **Stimulus** | **Phase** | **N** | **mean** | **sd** | **ci** | **mean** | **sd** | **ci** | **mean** | **sd** | **ci** | **mean** | **sd** | **ci** |
| Human | Isolation | 24 | 2.480 | 0.750 | 0.317 | 1.374 | 0.654 | 0.276 | 1.612 | 0.921 | 0.389 | 2.323 | 0.912 | 0.385 |
| Human | Reunion | 24 | 1.254 | 1.075 | 0.454 | 1.310 | 1.251 | 0.528 | 3.429 | 0.435 | 0.184 | 1.125 | 1.074 | 0.454 |
| Nothing | Isolation | 24 | 2.254 | 0.808 | 0.341 | 1.475 | 0.729 | 0.308 | 1.509 | 0.890 | 0.376 | 2.291 | 0.738 | 0.311 |
| Nothing | Reunion | 24 | 2.114 | 0.882 | 0.373 | 3.059 | 1.036 | 0.437 | 3.122 | 0.553 | 0.233 | 1.614 | 1.013 | 0.428 |
| Object | Isolation | 24 | 2.073 | 0.738 | 0.312 | 1.252 | 0.585 | 0.247 | 1.861 | 0.633 | 0.267 | 2.451 | 0.655 | 0.277 |
| Object | Reunion | 24 | 2.067 | 0.937 | 0.396 | 2.837 | 1.219 | 0.515 | 1.371 | 1.004 | 0.424 | 1.808 | 1.000 | 0.422 |
| **Day of Test** | | | | | | | | | | | | | | |
| Day1 | | 48 | 2.255 | 0.739 | 0.215 | 2.020 | 1.151 | 0.334 | 2.084 | 1.064 | 0.309 | 1.952 | 0.907 | 0.263 |
| Day2 | | 48 | 1.997 | 0.986 | 0.286 | 1.783 | 1.173 | 0.341 | 2.234 | 1.145 | 0.333 | 1.976 | 1.097 | 0.318 |
| Day3 | | 48 | 1.870 | 1.040 | 0.302 | 1.850 | 1.297 | 0.377 | 2.134 | 1.143 | 0.332 | 1.878 | 1.041 | 0.302 |

| | | | Number of zone changes | | | Time spent exploring the room (per min.) | | | Latency to enter proximal zone | | |
|---|---|---|---|---|---|---|---|---|---|---|---|
| **Stimulus** | **Phase** | **N** | **mean** | **sd** | **mean** | | **sd** | **ci** | **mean** | **sd** | **ci** |
| Human | Isolation | 24 | 2.079 | 0.476 | 0.201 | 2.831 | 0.502 | 0.212 | 229.568 | 103.919 | 43.881 |
| Human | Reunion | 24 | 1.941 | 0.451 | 0.190 | 2.410 | 0.569 | 0.240 | 195.342 | 279.449 | 118.001 |

| | | | | | | | | | | | |
|---|---|---|---|---|---|---|---|---|---|---|---|
| Nothing | Isolation | 24 | 2.059 | 0.516 | 0.218 | 2.805 | 0.683 | 0.289 | 272.457 | 77.145 | 32.575 |
| Nothing | Reunion | 24 | 1.937 | 0.389 | 0.164 | 2.267 | 0.651 | 0.275 | 248.653 | 273.280 | 115.396 |
| Object | Isolation | 24 | 2.227 | 0.318 | 0.134 | 3.195 | 0.520 | 0.220 | 255.447 | 91.274 | 38.542 |
| Object | Reunion | 24 | 1.923 | 0.632 | 0.267 | 2.677 | 0.791 | 0.334 | 555.383 | 117.831 | 49.756 |
| **Day of Test** | | | | | | | | | | | |
| Day1 | | 48 | 2.136 | 0.414 | 0.120 | 2.904 | 0.471 | 0.137 | 352.399 | 202.409 | 58.773 |
| Day2 | | 48 | 1.981 | 0.451 | 0.131 | 2.666 | 0.729 | 0.212 | 262.811 | 216.949 | 62.996 |
| Day3 | | 48 | 1.965 | 0.551 | 0.160 | 2.523 | 0.778 | 0.226 | 263.214 | 210.648 | 61.166 |

**Supplementary table S9**: Acoustic values for acoustic scores per significantly different groups (Stimulus, phase of test and location). The number of vocalisations per group is indicated and mean, standard deviation (sd), and 95% confidence interval (ci) are indicated for the vocalisation duration and all the spectral parameters used to build the spectral acoustic score (PC1)

| | | | **Mean (Hz)** | | | **Centroid (Hz)** | | | **Mean Dominant Frequency (KHz)** | | |
|---|---|---|---|---|---|---|---|---|---|---|---|
| **Stimulus** | **Phase of Test** | **N** | **mean** | **sd** | **ci** | **mean** | **sd** | **ci** | **mean** | **sd** | **ci** |
| Human | Isolation | 673 | 975.576 | 240.935 | 18.236 | 975.576 | 240.935 | 18.236 | 0.295 | 0.026 | 0.002 |
| Human | Reunion | 1302 | 1135.032 | 357.416 | 19.432 | 1135.032 | 357.416 | 19.432 | 0.307 | 0.065 | 0.004 |
| Nothing | Isolation | 775 | 1002.804 | 280.949 | 19.811 | 1002.804 | 280.949 | 19.811 | 0.295 | 0.037 | 0.003 |
| Nothing | Reunion | 1286 | 1018.402 | 307.527 | 16.824 | 1018.402 | 307.527 | 16.824 | 0.301 | 0.039 | 0.002 |
| Object | Isolation | 755 | 973.887 | 294.108 | 21.013 | 973.887 | 294.108 | 21.013 | 0.293 | 0.040 | 0.003 |
| Object | Reunion | 975 | 1042.705 | 292.903 | 18.408 | 1042.705 | 292.903 | 18.408 | 0.299 | 0.040 | 0.003 |
| **Location** | **Stimulus** | **N** | | | | | | | | | |

| | | | | | | | | | | | |
|---|---|---|---|---|---|---|---|---|---|---|---|
| Away | Human | 537 | 1157.254 | 382.488 | 32.424 | 1157.254 | 382.488 | 32.424 | 0.308 | 0.059 | 0.005 |
| Away | Object | 415 | 1034.390 | 303.160 | 29.253 | 1034.390 | 303.160 | 29.253 | 0.304 | 0.040 | 0.004 |
| Close | Human | 604 | 1127.452 | 337.411 | 26.963 | 1127.452 | 337.411 | 26.963 | 0.304 | 0.037 | 0.003 |
| Close | Object | 305 | 1074.428 | 257.501 | 29.014 | 1074.428 | 257.501 | 29.014 | 0.293 | 0.036 | 0.004 |
| | | | **Inter Quartile Range (Hz)** | | | **spectrum standard deviation** | | | **Call duration (s)** | | |
| **Stimulus** | **Phase of Test** | **N** | **mean** | **sd** | **ci** | **mean** | **sd** | **ci** | **mean** | **sd** | **ci** |
| Human | Isolation | 673 | 636.416 | 504.019 | 38.148 | 1473.751 | 254.950 | 111.544 | 0.379 | 0.222 | 0.017 |
| Human | Reunion | 1302 | 941.105 | 768.408 | 41.777 | 1603.690 | 296.579 | 87.190 | 0.251 | 0.185 | 0.010 |
| Nothing | Isolation | 775 | 719.745 | 617.101 | 43.514 | 1490.436 | 274.302 | 105.097 | 0.362 | 0.219 | 0.015 |
| Nothing | Reunion | 1286 | 749.850 | 675.886 | 36.975 | 1493.353 | 281.679 | 81.696 | 0.333 | 0.207 | 0.011 |
| Object | Isolation | 755 | 650.317 | 623.269 | 44.530 | 1461.763 | 274.854 | 104.436 | 0.385 | 0.241 | 0.017 |
| Object | Reunion | 975 | 776.602 | 633.862 | 39.836 | 1524.787 | 266.550 | 95.829 | 0.385 | 0.222 | 0.014 |
| **Location** | **Stimulus** | **N** | | | | | | | | | |
| Away | Human | 537 | 996.992 | 840.728 | 71.269 | 1617.188 | 313.709 | 137.089 | 0.288 | 0.196 | 0.017 |
| Away | Object | 415 | 755.197 | 653.837 | 63.091 | 1512.223 | 265.977 | 145.919 | 0.380 | 0.222 | 0.021 |
| Close | Human | 604 | 921.476 | 723.267 | 57.796 | 1600.669 | 287.511 | 127.910 | 0.200 | 0.145 | 0.012 |
| Close | Object | 305 | 835.885 | 560.337 | 63.136 | 1560.115 | 234.007 | 175.787 | 0.366 | 0.206 | 0.023 |
| | | | **Shannon entropy** | | | **Spectral Flatness** | | | **Entropy** | | |

| Stimulus | Phase of Test | N | mean | sd | ci | mean | sd | ci | mean | sd | ci |
|---|---|---|---|---|---|---|---|---|---|---|---|
| Human | Isolation | 673 | 0.651 | 0.067 | 0.005 | 0.270 | 0.087 | 0.007 | 0.501 | 0.049 | 0.004 |
| Human | Reunion | 1302 | 0.686 | 0.079 | 0.004 | 0.319 | 0.114 | 0.006 | 0.517 | 0.058 | 0.003 |
| Nothing | Isolation | 775 | 0.655 | 0.072 | 0.005 | 0.277 | 0.098 | 0.007 | 0.504 | 0.053 | 0.004 |
| Nothing | Reunion | 1286 | 0.660 | 0.077 | 0.004 | 0.280 | 0.104 | 0.006 | 0.506 | 0.058 | 0.003 |
| Object | Isolation | 755 | 0.647 | 0.074 | 0.005 | 0.267 | 0.100 | 0.007 | 0.498 | 0.054 | 0.004 |
| Object | Reunion | 975 | 0.669 | 0.074 | 0.005 | 0.291 | 0.098 | 0.006 | 0.516 | 0.054 | 0.003 |
| **Location** | **Stimulus** | **N** | | | | | | | | | |
| Away | Human | 537 | 0.686 | 0.085 | 0.007 | 0.322 | 0.119 | 0.010 | 0.521 | 0.062 | 0.005 |
| Away | Object | 415 | 0.665 | 0.072 | 0.007 | 0.285 | 0.098 | 0.009 | 0.512 | 0.053 | 0.005 |
| Close | Human | 604 | 0.689 | 0.076 | 0.006 | 0.320 | 0.112 | 0.009 | 0.515 | 0.056 | 0.004 |
| Close | Object | 305 | 0.685 | 0.068 | 0.008 | 0.306 | 0.089 | 0.010 | 0.527 | 0.049 | 0.005 |

**Supplementary table S10**: After model validation, effects of explanatory variables were computed using the ‘Anova’ function (‘car’ R package), running Type II Wald chisquare trials. All linear models were computed using the ’lmer’ function, taking into account repeated observations as random factors (individual). One model had a binary response variable and one model was on counting was computed with a generalized model taking into account repeated observations as random factors, respectively using a Binomial and a Poisson distribution (see models 2 and 4 when indicated). P values were considered significant when below 0.05.

| | Chisq | Df | Pr.Chisq. |
|---|---|---|---|
| **Model 1: PC1 of choice test** | | | |
| Stimulus | 0.284 | 1 | 0.594 |
| Day | 0.086 | 1 | 0.769 |
| Position of human (left vs. right) | 0.073 | 1 | 0.787 |
| Stimulus: Day | 6.300 | 1 | 0.012 |
| **Model 1: PC2 of choice test** | | | |
| Stimulus | 7.286 | 1 | 0.007 |
| Day | 3.252 | 1 | 0.071 |
| Position of human (left vs. right) | 0.166 | 1 | 0.683 |
| Stimulus: Day | 0.715 | 1 | 0.398 |
| **Model 1: PC3 of choice test** | | | |
| Stimulus | 1.512 | 1 | 0.219 |
| Day | 0.567 | 1 | 0.451 |
| Position of human (left vs. right) | 0.019 | 1 | 0.891 |
| Stimulus: Day | 1.973 | 1 | 0.160 |
| **Model 2 : first approach (binomial)** | | | |
| Day | 3.440 | 1 | 0.064 |
| Position of human (left vs. right) | 1.831 | 1 | 0.176 |
| **Model 3 : Isolation/Reunion test - behaviour PC1** | | | |
| Stimulus | 24.063 | 2 | <0.001 |
| Phase of Test | 33.834 | 1 | <0.001 |
| Day | 10.071 | 2 | 0.007 |
| Stimulus : Phase of Test | 16.565 | 2 | <0.001 |
| Stimulus : Day | 3.035 | 4 | 0.552 |

| | | | |
|---|---|---|---|
| Phase of Test : Day | 1.474 | 2 | 0.479 |
| **Model 3 : Isolation/Reunion test - behaviour PC2** | | | |
| Stimulus | 16.576 | 2 | <0.001 |
| Phase of Test | 45.999 | 1 | <0.001 |
| Day | 0.994 | 2 | 0.608 |
| Stimulus : Phase of Test | 41.531 | 2 | <0.001 |
| Stimulus : Day | 4.211 | 4 | 0.378 |
| Phase of Test : Day | 4.887 | 2 | 0.087 |
| **Model 3 : Isolation/Reunion test - behaviour PC3** | | | |
| Stimulus | 29.694 | 2 | <0.001 |
| Phase of Test | 44.445 | 1 | <0.001 |
| Day | 0.201 | 2 | 0.904 |
| Stimulus : Phase of Test | 36.383 | 2 | <0.001 |
| Stimulus : Day | 7.294 | 4 | 0.121 |
| Phase of Test : Day | 2.922 | 2 | 0.232 |
| **Model 3 : Acoustic Spectral Score (PC1)** | | | |
| Stimulus | 46.813 | 2 | <0.001 |
| Phase of Test | 69.814 | 1 | <0.001 |
| Day | 12.796 | 2 | 0.002 |
| Stimulus : Phase of Test | 45.131 | 2 | <0.001 |
| Stimulus : Day | 26.773 | 4 | <0.001 |
| Phase of Test : Day | 1.301 | 2 | 0.522 |
| **Model 3 : Acoustic grunt duration (log)** | | | |
| Stimulus | 257.550 | 2 | <0.001 |
| Phase of Test | 129.889 | 1 | <0.001 |
| Day | 19.919 | 2 | <0.001 |
| Stimulus : Phase of Test | 210.139 | 2 | <0.001 |
| Stimulus : Day | 4.579 | 4 | 0.333 |
| Phase of Test : Day | 4.761 | 2 | 0.093 |

**Supplementary table S11**: Post hoc tests on models following significant interactions or single effects of explanatory variables. Each model is indicated, all post hoc tests were computed using the 'lsmeans' function with Tukey correction for multiple testing. P values were considered significant when below 0.05.

| Contrast | Estimate | SE | DF | t.ratio | p.value |
|---|---|---|---|---|---|
| **Model 1: PC1 of choice test** | | | | | |
| Human, day1 - Object, day1 | -0.716 | 0.512 | 68 | -1.398 | 0.505 |
| Human, day1 - Human, day2 | -0.803 | 0.513 | 68 | -1.563 | 0.406 |
| Human, day1 - Object, day2 | 0.300 | 0.513 | 68 | 0.584 | 0.937 |
| Object, day1 - Human, day2 | -0.086 | 0.513 | 68 | -0.168 | 0.998 |
| Object, day1 - Object, day2 | 1.016 | 0.513 | 68 | 1.980 | 0.206 |
| Human, day2 - Object, day2 | 1.102 | 0.512 | 68 | 2.151 | 0.148 |
| **Model 1: PC2 of choice test** | | | | | |
| Human - Object | 0.475 | 0.176 | 68 | 2.699 | 0.009 |
| Day1 - Day2 | 0.318 | 0.177 | 68 | 1.803 | 0.076 |
| **Model 3 : Isolation/Reunion test - behaviour PC1** | | | | | |
| Human,Isolation - Object,Isolation | -0.041 | 0.330 | 109 | -0.125 | 1.000 |
| Human,Isolation - Nothing,Isolation | 0.192 | 0.330 | 109 | 0.582 | 0.992 |
| Human,Isolation - Human,Reunion | -2.088 | 0.330 | 109 | -6.332 | <0.001 |
| Human,Isolation - Object,Reunion | -1.082 | 0.330 | 109 | -3.282 | <0.001 |
| Human,Isolation - Nothing,Reunion | -0.001 | 0.330 | 109 | -0.004 | 1.000 |
| Object,Isolation - Nothing,Isolation | 0.233 | 0.330 | 109 | 0.708 | 0.981 |
| Object,Isolation - Human,Reunion | -2.047 | 0.330 | 109 | -6.207 | <0.001 |
| Object,Isolation - Object,Reunion | -1.041 | 0.330 | 109 | -3.156 | 0.025 |
| Object,Isolation - Nothing,Reunion | 0.040 | 0.330 | 109 | 0.121 | 1.000 |
| Nothing,Isolation - Human,Reunion | -2.280 | 0.330 | 109 | -6.914 | <0.001 |
| Nothing,Isolation - Object,Reunion | -1.274 | 0.330 | 109 | -3.864 | 0.003 |
| Nothing,Isolation - Nothing,Reunion | -0.193 | 0.330 | 109 | -0.587 | 0.992 |
| Human,Reunion - Object,Reunion | 1.006 | 0.330 | 109 | 3.050 | 0.033 |
| Human,Reunion - Nothing,Reunion | 2.087 | 0.330 | 109 | 6.328 | <0.001 |
| Object,Reunion - Nothing,Reunion | 1.081 | 0.330 | 109 | 3.277 | 0.017 |

**Model 3 : Isolation/Reunion test - behaviour PC1**

| | | | | | |
|---|---|---|---|---|---|
| Day1 - Day2 | -0.552 | 0.233 | 109 | -2.366 | 0.051 |
| Day1 - Day3 | -0.703 | 0.233 | 109 | -3.015 | 0.009 |
| Day2 - Day3 | -0.151 | 0.233 | 109 | -0.648 | 0.794 |

**Model 3 : Isolation/Reunion test - behaviour PC2**

| | | | | | |
|---|---|---|---|---|---|
| Human,Isolation - Object,Isolation | 0.115 | 0.258 | 109 | 0.443 | 0.998 |
| Human,Isolation - Nothing,Isolation | -0.586 | 0.258 | 109 | -2.269 | 0.216 |
| Human,Isolation - Human,Reunion | -0.154 | 0.258 | 109 | -0.598 | 0.991 |
| Human,Isolation - Object,Reunion | 1.104 | 0.258 | 109 | 4.273 | 0.001 |
| Human,Isolation - Nothing,Reunion | 1.614 | 0.258 | 109 | 6.246 | <0.001 |
| Object,Isolation - Nothing,Isolation | -0.701 | 0.258 | 109 | -2.712 | 0.081 |
| Object,Isolation - Human,Reunion | -0.269 | 0.258 | 109 | -1.041 | 0.903 |
| Object,Isolation - Object,Reunion | 0.990 | 0.258 | 109 | 3.830 | 0.003 |
| Object,Isolation - Nothing,Reunion | 1.500 | 0.258 | 109 | 5.803 | <0.001 |
| Nothing,Isolation - Human,Reunion | 0.432 | 0.258 | 109 | 1.671 | 0.554 |
| Nothing,Isolation - Object,Reunion | 1.691 | 0.258 | 109 | 6.542 | <0.001 |
| Nothing,Isolation - Nothing,Reunion | 2.200 | 0.258 | 109 | 8.515 | <0.001 |
| Human,Reunion - Object,Reunion | 1.259 | 0.258 | 109 | 4.871 | <0.001 |
| Human,Reunion - Nothing,Reunion | 1.769 | 0.258 | 109 | 6.844 | <0.001 |
| Object,Reunion - Nothing,Reunion | 0.510 | 0.258 | 109 | 1.973 | 0.365 |

**Model 3 : Isolation/Reunion test - behaviour PC3**

| | | | | | |
|---|---|---|---|---|---|
| Human,Isolation - Object,Isolation | -0.150 | 0.210 | 109 | -0.713 | 0.980 |
| Human,Isolation - Nothing,Isolation | -0.032 | 0.210 | 109 | -0.153 | 1.000 |
| Human,Isolation - Human,Reunion | 1.002 | 0.210 | 109 | 4.760 | <0.001 |
| Human,Isolation - Object,Reunion | 1.446 | 0.210 | 109 | 6.872 | <0.001 |
| Human,Isolation - Nothing,Reunion | -0.200 | 0.210 | 109 | -0.951 | 0.932 |
| Object,Isolation - Nothing,Isolation | 0.118 | 0.210 | 109 | 0.560 | 0.993 |
| Object,Isolation - Human,Reunion | 1.152 | 0.210 | 109 | 5.473 | <0.001 |
| Object,Isolation - Object,Reunion | 1.596 | 0.210 | 109 | 7.585 | <0.001 |

| | | | | | |
|---|---|---|---|---|---|
| Object,Isolation - Nothing,Reunion | -0.050 | 0.210 | 109 | -0.238 | 1.000 |
| Nothing,Isolation - Human,Reunion | 1.034 | 0.210 | 109 | 4.913 | <0.001 |
| Nothing,Isolation - Object,Reunion | 1.478 | 0.210 | 109 | 7.025 | <0.001 |
| Nothing,Isolation - Nothing,Reunion | -0.168 | 0.210 | 109 | -0.798 | 0.967 |
| Human,Reunion - Object,Reunion | 0.444 | 0.210 | 109 | 2.112 | 0.289 |
| Human,Reunion - Nothing,Reunion | -1.202 | 0.210 | 109 | -5.711 | <0.001 |
| Object,Reunion - Nothing,Reunion | -1.646 | 0.210 | 109 | -7.823 | <0.001 |
| **Model 3 : Acoustic Spectral Score (PC1)** | | | | | |
| Human,isolation - Nothing,isolation | 0.177 | 0.129 | 5735 | 1.377 | 0.741 |
| Human,isolation - Object,isolation | -0.056 | 0.131 | 5737 | -0.427 | 0.998 |
| Nothing,isolation - Object,isolation | -0.233 | 0.126 | 5736 | -1.854 | 0.431 |
| Human,isolation - Human,reunion | 1.088 | 0.117 | 5743 | 9.289 | <0.001 |
| Human,isolation - Nothing,reunion | 0.182 | 0.117 | 5737 | 1.550 | 0.632 |
| Human,isolation - Object,reunion | 0.574 | 0.123 | 5737 | 4.660 | <0.001 |
| Nothing,isolation - Human,reunion | 0.911 | 0.112 | 5741 | 8.163 | <0.001 |
| Nothing,isolation - Nothing,reunion | 0.004 | 0.112 | 5738 | 0.038 | 1.000 |
| Nothing,isolation - Object,reunion | 0.396 | 0.118 | 5741 | 3.346 | 0.011 |
| Object,isolation - Human,reunion | 1.144 | 0.114 | 5746 | 10.012 | <0.001 |
| Object,isolation - Nothing,reunion | 0.238 | 0.113 | 5739 | 2.096 | 0.289 |
| Object,isolation - Object,reunion | 0.629 | 0.120 | 5740 | 5.251 | <0.001 |
| Human,reunion - Nothing,reunion | -0.907 | 0.098 | 5747 | -9.223 | <0.001 |
| Human,reunion - Object,reunion | -0.515 | 0.105 | 5745 | -4.880 | <0.001 |
| Nothing,reunion - Object,reunion | 0.392 | 0.105 | 5744 | 3.725 | 0.003 |
| **Model 3 : Acoustic Spectral Score (PC1)** | | | | | |
| Human, Day1 - Nothing, Day1 | -0.025 | 0.464 | 24 | -0.055 | 1.000 |
| Human, Day1 - Object, Day1 | -0.055 | 0.464 | 24 | -0.119 | 1.000 |
| Human, Day1 - Human, Day2 | 0.631 | 0.463 | 23 | 1.364 | 0.900 |
| Human, Day1 - Nothing, Day2 | 0.117 | 0.136 | 5736 | 0.856 | 0.995 |
| Human, Day1 - Object, Day2 | -0.139 | 0.468 | 24 | -0.298 | 1.000 |

| | | | | | |
|---|---|---|---|---|---|
| Human, Day1 - Human, Day3 | 0.361 | 0.466 | 24 | 0.775 | 0.997 |
| Human, Day1 - Nothing, Day3 | -0.193 | 0.465 | 24 | -0.414 | 1.000 |
| Human, Day1 - Object, Day3 | 0.331 | 0.147 | 5748 | 2.259 | 0.368 |
| Nothing, Day1 - Object, Day1 | -0.030 | 0.465 | 24 | -0.064 | 1.000 |
| Nothing, Day1 - Human, Day2 | 0.656 | 0.464 | 24 | 1.414 | 0.881 |
| Nothing, Day1 - Nothing, Day2 | 0.142 | 0.462 | 23 | 0.307 | 1.000 |
| Nothing, Day1 - Object, Day2 | -0.114 | 0.151 | 5734 | -0.755 | 0.998 |
| Nothing, Day1 - Human, Day3 | 0.387 | 0.146 | 5741 | 2.656 | 0.164 |
| Nothing, Day1 - Nothing, Day3 | -0.167 | 0.466 | 24 | -0.359 | 1.000 |
| Nothing, Day1 - Object, Day3 | 0.356 | 0.465 | 24 | 0.766 | 0.997 |
| Object, Day1 - Human, Day2 | 0.686 | 0.132 | 5738 | 5.181 | 0.000 |
| Object, Day1 - Nothing, Day2 | 0.172 | 0.461 | 23 | 0.373 | 1.000 |
| Object, Day1 - Object, Day2 | -0.084 | 0.467 | 24 | -0.180 | 1.000 |
| Object, Day1 - Human, Day3 | 0.416 | 0.467 | 24 | 0.892 | 0.991 |
| Object, Day1 - Nothing, Day3 | -0.137 | 0.138 | 5743 | -0.996 | 0.986 |
| Object, Day1 - Object, Day3 | 0.386 | 0.464 | 24 | 0.832 | 0.995 |
| Human, Day2 - Nothing, Day2 | -0.514 | 0.460 | 23 | -1.119 | 0.965 |
| Human, Day2 - Object, Day2 | -0.770 | 0.466 | 24 | -1.651 | 0.768 |
| Human, Day2 - Human, Day3 | -0.270 | 0.466 | 24 | -0.579 | 1.000 |
| Human, Day2 - Nothing, Day3 | -0.824 | 0.133 | 5742 | -6.171 | 0.000 |
| Human, Day2 - Object, Day3 | -0.300 | 0.463 | 24 | -0.647 | 0.999 |
| Nothing, Day2 - Object, Day2 | -0.256 | 0.465 | 24 | -0.551 | 1.000 |
| Nothing, Day2 - Human, Day3 | 0.245 | 0.464 | 24 | 0.527 | 1.000 |
| Nothing, Day2 - Nothing, Day3 | -0.309 | 0.462 | 23 | -0.669 | 0.999 |
| Nothing, Day2 - Object, Day3 | 0.214 | 0.135 | 5744 | 1.584 | 0.814 |
| Object, Day2 - Human, Day3 | 0.501 | 0.154 | 5738 | 3.253 | 0.032 |
| Object, Day2 - Nothing, Day3 | -0.053 | 0.469 | 25 | -0.114 | 1.000 |
| Object, Day2 - Object, Day3 | 0.470 | 0.468 | 25 | 1.004 | 0.982 |
| Human, Day3 - Nothing, Day3 | -0.554 | 0.468 | 24 | -1.184 | 0.953 |

| | | | | | |
|---|---|---|---|---|---|
| Human, Day3 - Object, Day3 | -0.030 | 0.467 | 24 | -0.065 | 1.000 |
| Nothing, Day3 - Object, Day3 | 0.524 | 0.466 | 24 | 1.125 | 0.964 |
| **Model 3 : Acoustic grunt duration (log)** | | | | | |
| Human,isolation - Nothing,isolation | 0.080 | 0.031 | 5734 | 2.604 | 0.096 |
| Human,isolation - Object,isolation | 0.029 | 0.031 | 5736 | 0.943 | 0.935 |
| Nothing,isolation - Object,isolation | -0.051 | 0.030 | 5735 | -1.685 | 0.542 |
| Human,isolation - Human,reunion | 0.513 | 0.028 | 5741 | 18.340 | <0.001 |
| Human,isolation - Nothing,reunion | 0.122 | 0.028 | 5735 | 4.353 | <0.001 |
| Human,isolation - Object,reunion | 0.023 | 0.029 | 5736 | 0.795 | 0.969 |
| Nothing,isolation - Human,reunion | 0.433 | 0.027 | 5739 | 16.250 | <0.001 |
| Nothing,isolation - Nothing,reunion | 0.042 | 0.027 | 5736 | 1.560 | 0.625 |
| Nothing,isolation - Object,reunion | -0.057 | 0.028 | 5739 | -2.008 | 0.338 |
| Object,isolation - Human,reunion | 0.483 | 0.027 | 5744 | 17.721 | <0.001 |
| Object,isolation - Nothing,reunion | 0.092 | 0.027 | 5737 | 3.413 | 0.008 |
| Object,isolation - Object,reunion | -0.006 | 0.029 | 5738 | -0.214 | 1.000 |
| Human,reunion - Nothing,reunion | -0.391 | 0.023 | 5745 | -16.665 | <0.001 |
| Human,reunion - Object,reunion | -0.489 | 0.025 | 5743 | -19.448 | <0.001 |
| Nothing,reunion - Object,reunion | -0.098 | 0.025 | 5742 | -3.921 | 0.001 |
| **Model 3 : Acoustic grunt duration (log)** | | | | | |
| Day1 - Day2 | 0.076 | 0.020 | 5734 | 3.892 | <0.001 |
| Day1 - Day3 | 0.052 | 0.020 | 5742 | 2.604 | 0.025 |
| Day2 - Day3 | -0.024 | 0.020 | 5739 | -1.240 | 0.430 |

**Supplementary table S12**: Table of model estimates, following significant effect of explanatory variables. All estimates were calculated using the 'lsmeans' function ('lmerTest' R package) and are presented per relevant group lsmean, SE, DF, lower.CI upper.CI respectively represent mean estimates, standard error, degrees of freedom and lower and upper limit of 95% confidence interval

| Factor 1 | Factor 2 | Lsmean | SE | DF | Lower.CL | Upper.CL |
|---|---|---|---|---|---|---|
| **Model 1: PC1 of choice test** | | | | | | |
| Human | Day1 | -0.305 | 0.363 | 90.995 | -1.025 | 0.416 |
| Object | Day1 | 0.412 | 0.363 | 90.995 | -0.309 | 1.132 |
| Human | Day2 | 0.498 | 0.363 | 90.995 | -0.223 | 1.218 |
| Object | Day2 | -0.605 | 0.363 | 90.995 | -1.325 | 0.116 |
| **Model 1: PC2 of choice test** | | | | | | |
| Human | - | 0.238 | 0.200 | 34.646 | -0.169 | 0.644 |
| Object | - | -0.238 | 0.200 | 34.646 | -0.644 | 0.169 |
| - | Day1 | 0.159 | 0.200 | 34.731 | -0.248 | 0.566 |
| - | Day2 | -0.159 | 0.200 | 34.731 | -0.566 | 0.248 |
| **Model 2 : first approach (binomial)** | | | | | | |
| Day1 | - | -0.211 | 0.421 | Inf | -1.036 | 0.613 |
| Day2 | - | 0.961 | 0.465 | Inf | 0.051 | 1.872 |
| **Model 3 : Isolation/Reunion test - behaviour PC1** | | | | | | |
| Human | Isolation | -0.406 | 0.291 | 117.573 | -0.982 | 0.170 |
| Object | Isolation | -0.329 | 0.291 | 117.573 | -0.905 | 0.247 |
| Nothing | Isolation | -0.686 | 0.291 | 117.573 | -1.262 | -0.110 |
| Human | Reunion | 1.277 | 0.291 | 117.573 | 0.701 | 1.853 |
| Object | Reunion | 0.271 | 0.291 | 117.573 | -0.305 | 0.848 |
| Nothing | Reunion | -0.128 | 0.291 | 117.573 | -0.704 | 0.448 |
| **Model 3 : Isolation/Reunion test - behaviour PC1** | | | | | | |
| Day1 | - | -0.473 | 0.219 | 71.500 | -0.910 | -0.036 |
| Day2 | - | 0.124 | 0.219 | 71.500 | -0.313 | 0.562 |
| Day3 | - | 0.349 | 0.219 | 71.500 | -0.089 | 0.786 |
| **Model 3 : Isolation/Reunion test - behaviour PC2** | | | | | | |

| | | | | | | |
|---|---|---|---|---|---|---|
| Human | Isolation | 0.031 | 0.202 | 121.263 | -0.370 | 0.432 |
| Object | Isolation | -0.154 | 0.202 | 121.263 | -0.555 | 0.247 |
| Nothing | Isolation | 0.490 | 0.202 | 121.263 | 0.089 | 0.891 |
| Human | Reunion | 1.095 | 0.202 | 121.263 | 0.694 | 1.496 |
| Object | Reunion | -0.047 | 0.202 | 121.263 | -0.448 | 0.354 |
| Nothing | Reunion | -1.415 | 0.202 | 121.263 | -1.816 | -1.014 |
| **Model 3 : Isolation/Reunion test - behaviour PC3** | | | | | | |
| Human | Isolation | -0.496 | 0.177 | 91.865 | -0.848 | -0.144 |
| Object | Isolation | -0.543 | 0.177 | 91.865 | -0.895 | -0.191 |
| Nothing | Isolation | -0.605 | 0.177 | 91.865 | -0.957 | -0.253 |
| Human | Reunion | 0.390 | 0.177 | 91.865 | 0.038 | 0.742 |
| Object | Reunion | 1.262 | 0.177 | 91.865 | 0.910 | 1.614 |
| Nothing | Reunion | -0.009 | 0.177 | 91.865 | -0.361 | 0.343 |
| **Model 3 : Acoustic Spectral Score (PC1)** | | | | | | |
| Human | isolation | 0.319 | 0.204 | 31.821 | -0.096 | 0.735 |
| Nothing | isolation | 0.142 | 0.201 | 30.212 | -0.269 | 0.552 |
| Object | isolation | 0.375 | 0.202 | 30.677 | -0.037 | 0.787 |
| Human | reunion | -0.769 | 0.194 | 26.023 | -1.164 | -0.374 |
| Nothing | reunion | 0.138 | 0.194 | 25.830 | -0.257 | 0.532 |
| Object | reunion | -0.254 | 0.197 | 27.835 | -0.656 | 0.148 |
| **Model 3 : Acoustic Spectral Score (PC1)** | | | | | | |
| Human | Day1 | 0.106 | 0.328 | 23.664 | -0.572 | 0.783 |
| Nothing | Day1 | 0.131 | 0.330 | 24.068 | -0.549 | 0.812 |
| Object | Day1 | 0.161 | 0.328 | 23.596 | -0.516 | 0.838 |
| Human | Day2 | -0.525 | 0.327 | 23.254 | -1.200 | 0.150 |
| Nothing | Day2 | -0.011 | 0.324 | 22.550 | -0.680 | 0.659 |
| Object | Day2 | 0.245 | 0.333 | 25.181 | -0.443 | 0.933 |
| Human | Day3 | -0.255 | 0.332 | 24.806 | -0.941 | 0.431 |
| Nothing | Day3 | 0.299 | 0.330 | 24.084 | -0.382 | 0.980 |

| | | | | | | |
|---|---|---|---|---|---|---|
| Object | Day3 | -0.225 | 0.329 | 23.838 | -0.904 | 0.454 |
| **Model 3 : Acoustic grunt duration (log)** | | | | | | |
| Human | isolation | -1.153 | 0.054 | 29.113 | -1.263 | -1.042 |
| Nothing | isolation | -1.233 | 0.053 | 27.912 | -1.342 | -1.124 |
| Object | isolation | -1.182 | 0.054 | 28.262 | -1.292 | -1.073 |
| Human | reunion | -1.665 | 0.052 | 24.753 | -1.771 | -1.559 |
| Nothing | reunion | -1.274 | 0.052 | 24.605 | -1.380 | -1.169 |
| Object | reunion | -1.176 | 0.052 | 26.121 | -1.283 | -1.069 |
| **Model 3 : Acoustic grunt duration (log)** | | | | | | |
| Day1 | - | -1.238 | 0.051 | 23.156 | -1.343 | -1.133 |
| Day2 | - | -1.314 | 0.051 | 23.035 | -1.419 | -1.209 |
| Day3 | - | -1.290 | 0.051 | 23.406 | -1.395 | -1.184 |

**Supplementary table S13**: Model selection table. Only the equivalent best models after selections are shown (i.e Delta AICc<2) as well as the null model. All models are ranked. Estimates of each covariate when present in a given model are given, a '+' means the term is present in the model, DF, LogLik, AICc, Delta AICc and weight of models. 'Stim', 'Loc.' and 'Behav. Prox.' refer to 'Stimulus', 'Location' and 'behavioural proximity'. When the null model is in the best selected model, then we cannot extract any predictor.

| Response variable | rank | Intercept | Stim | Day | Loc. | Int. Index | - Behav. Prox. | Stim :Day | Stim :Loc. | Stim :Int. Index | Stim : - Behav. Prox. | Loc : - Behav. Prox. | df | logLik | AICc | delta | weight |
|---|---|---|---|---|---|---|---|---|---|---|---|---|---|---|---|---|---|
| Total Number grunts (Poisson) | 1 | 2.77 | + | + | + | | 0.03 | | + | | + | | 9 | -487.236 | 994.751 | 0.000 | 0.426 |
| | 2 | 2.77 | + | + | + | | 0.07 | | + | | + | + | 10 | -486.094 | 995.008 | 0.257 | 0.375 |
| | 66 | 2.73 | NULL | | | | | | | | | | 2 | -581.376 | 1166.891 | 172.140 | 0.000 |
| Grunt rate (log) | 1 | -1.56 | + | | | | | | | | | | 4 | -176.946 | 362.117 | 0.000 | 0.193 |
| | 2 | -1.56 | + | | | | 0.04 | | | | | | 5 | -176.716 | 363.774 | 1.656 | 0.084 |
| | 119 | -1.77 | NULL | | | | | | | | | | 3 | -187.641 | 381.416 | 19.299 | 0.000 |
| Grunt duration (log) | 1 | -1.50 | + | + | + | 0.11 | -0.05 | | + | + | | | 11 | -1600.120 | 3222.382 | 0.000 | 0.172 |
| | 2 | -1.49 | + | + | + | 0.10 | -0.05 | | + | | | | 10 | -1601.159 | 3222.436 | 0.054 | 0.167 |
| | 3 | -1.50 | + | + | + | 0.12 | -0.06 | | + | + | | + | 12 | -1599.841 | 3223.851 | 1.470 | 0.082 |
| | 4 | -1.50 | + | + | + | 0.11 | -0.06 | | + | + | + | | 12 | -1599.848 | 3223.866 | 1.484 | 0.082 |
| | 5 | -1.49 | + | + | + | 0.10 | -0.06 | | + | | | | 11 | -1600.872 | 3223.887 | 1.505 | 0.081 |

| Response variable | rank | Intercept | Stim | Day | Loc. | Int. Index | -Behav. Prox. 1 | Stim :Day | Stim :Loc. | Stim :Int. Index | Stim : -Behav. Prox. 1 | Loc : -Behav. Prox. 1 | df | logLik | AICc | delta | weight |
|---|---|---|---|---|---|---|---|---|---|---|---|---|---|---|---|---|---|
| | 6 | -1.49 | + | + | + | 0.10 | -0.06 | | + | | + | | 11 | -1600.948 | 3224.039 | 1.657 | 0.075 |
| | 137 | -1.49 | NULL | | | | | | | | | | 3 | -1814.290 | 3634.593 | 412.211 | 0.000 |
| Acoustic spectral score (PC1) | 1 | -0.18 | + | + | + | 0.27 | -0.28 | | + | | | | 10 | -4318.392 | 8656.904 | 0.000 | 0.101 |
| | 2 | -0.15 | + | + | + | 0.28 | -0.16 | | + | | + | | 11 | -4317.445 | 8657.032 | 0.128 | 0.095 |
| | 3 | -0.31 | + | + | + | 0.28 | -0.28 | + | + | | | | 12 | -4316.490 | 8657.149 | 0.245 | 0.089 |
| | 4 | -0.30 | + | + | + | 0.28 | -0.17 | + | + | | + | | 13 | -4315.669 | 8657.534 | 0.630 | 0.074 |
| | 5 | -0.20 | + | + | + | 0.31 | -0.28 | | + | + | | | 11 | -4318.078 | 8658.298 | 1.394 | 0.050 |
| | 6 | -0.17 | + | + | + | 0.30 | -0.17 | | + | + | + | | 12 | -4317.192 | 8658.553 | 1.649 | 0.044 |
| | 7 | -0.33 | + | + | + | 0.31 | -0.28 | + | + | + | | | 13 | -4316.190 | 8658.578 | 1.674 | 0.044 |
| | 8 | -0.15 | + | + | + | 0.27 | -0.13 | | + | | + | + | 12 | -4317.346 | 8658.860 | 1.956 | 0.038 |
| | 9 | -0.19 | + | + | + | 0.27 | -0.27 | | + | | | | 11 | -4318.362 | 8658.866 | 1.962 | 0.038 |

| | | | | | | | | | | | | | | | | | |
|---|---|---|---|---|---|---|---|---|---|---|---|---|---|---|---|---|---|
| | 137 | -0.38 | NULL | | | | | | | | | | 3 | -4346.453 | 8698.918 | 42.014 | 0.000 |

| **Response variable** | **rank** | **Intercept** | **Stim** | **Day** | **Loc.** | **Int. Index** | **-Behav. Prox. 1** | **Stim :Day** | **Stim :Loc.** | **Stim :Int. Index** | **Stim : -Behav. Prox. 1** | **Loc : -Behav. Prox. 1** | **df** | **logLik** | **AICc** | **delta** | **weight** |
|---|---|---|---|---|---|---|---|---|---|---|---|---|---|---|---|---|---|
| **More than 1 grunt per interval *(Binomial)*** | 1 | 1.49 | + | | | -0.21 | | | | | | | 4 | -153.427 | 314.996 | 0.000 | 0.071 |
| | 2 | 1.46 | + | | | | | | | | | | 3 | -154.502 | 315.090 | 0.094 | 0.068 |
| | 3 | 1.77 | + | + | | | | | | | | | 5 | -153.031 | 316.277 | 1.282 | 0.038 |
| | 4 | 1.63 | + | | + | -0.22 | | | | | | | 5 | -153.131 | 316.476 | 1.480 | 0.034 |
| | 5 | 1.79 | + | + | | -0.20 | | | | | | | 6 | -152.124 | 316.550 | 1.554 | 0.033 |
| | 6 | 1.56 | + | | + | | | | | | | | 4 | -154.291 | 316.725 | 1.730 | 0.030 |
| | 7 | 1.19 | NULL | | | | | | | | | | 2 | -156.355 | 316.752 | 1.756 | 0.030 |
| | 8 | 1.50 | + | | | -0.22 | 0.08 | | | | | | 5 | -153.319 | 316.853 | 1.857 | 0.028 |
| | 9 | 1.46 | + | | | | 0.08 | | | | | | 4 | -154.410 | 316.963 | 1.967 | 0.027 |

### 13 Supplementary figures

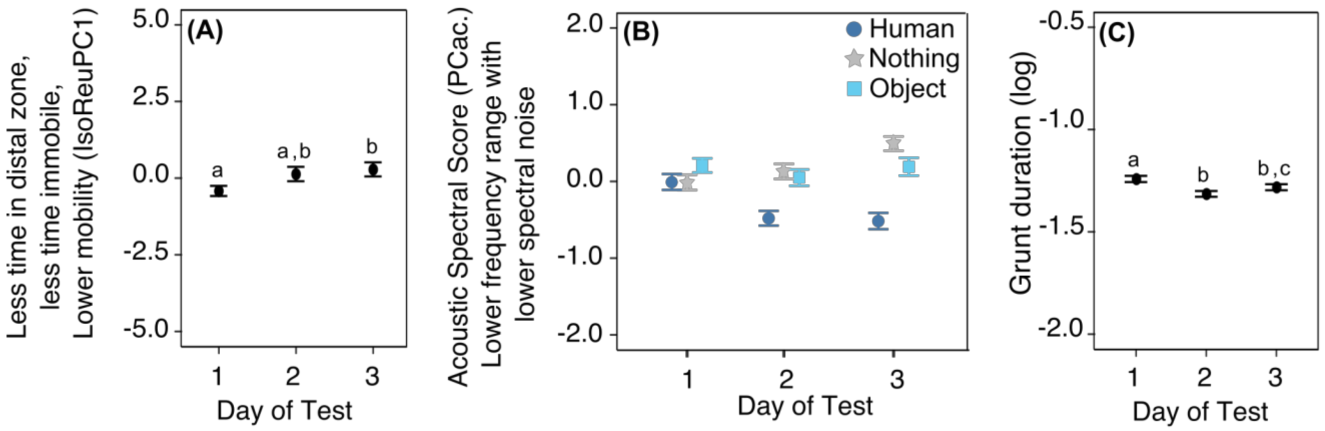


**Supplementary Figure S5**: Effect of the day of the test on vocal and spatial behavior of piglets during isolation/reunion tests. Mean +- se of the first behavioral score (A), the acoustic spectral score of grunts (B) and grunt duration (C), according to the day and/or the type of reunion (B, dark blue circles for human, light blue square for object and grey stars for nothing). Different letters show significantly different groups ($p < 0.05$). All model estimates, anova tables and results of post hoc tests are available in supplementary tables S4-S6.
